# Tonic endocannabinoid signaling supports sleep through development in both sexes

**DOI:** 10.1101/2021.05.10.443432

**Authors:** Sheneé C. Martin, Sean M. Gay, Michael L. Armstrong, Nila M. Pazhayam, Nichole Reisdorph, Graham H. Diering

## Abstract

Sleep is an essential behavior that supports brain function and cognition throughout life, in part by acting on neuronal synapses. The synaptic signaling pathways that mediate the restorative benefits of sleep are not fully understood, particularly in the context of development. Endocannabinoids (eCBs) including 2-arachidonyl glycerol (2-AG) and anandamide (AEA), are bioactive lipids that activate cannabinoid receptor, CB1, to regulate synaptic transmission and mediate cognitive functions and many behaviors, including sleep. We used targeted mass spectrometry to measure changes in forebrain synaptic eCBs during the sleep/wake cycle in developing and adult mice. We find that eCBs are downregulated in response to acute sleep deprivation in juvenile mice, while in young adults eCBs are upregulated during the sleep phase in a circadian manner. Next we manipulated the eCB system using selective pharmacology and measured the effects on sleep behavior in developing and adult mice of both sexes using a non-invasive piezoelectric home-cage recording apparatus. Enhancement of eCB signaling through inhibition of 2-AG or AEA degradation, increased dark phase sleep amount and bout length in developing and adult males, but not in females. Inhibition of CB1 by injection of the antagonist AM251 reduced sleep time and caused sleep fragmentation in developing and adult males and females. Our data suggest that males are more sensitive to the sleep promoting effects of enhanced eCBs but that tonic eCB signaling supports sleep behavior through multiple stages of development in both sexes. This work informs the further development of cannabinoid-based therapeutics for sleep disruption.

## Introduction

Sleep is an essential conserved behavior seen throughout life. Chronic sleep disruption is associated with many diseases ranging from neurodevelopmental disorders in earlier life to neurodegenerative diseases later in life. Indeed, sleep disruption is a major comorbidity in autism spectrum disorder (ASD), reported in up to 80% of affected individuals (Mazzone et al., 2018; Missig et al., 2019). Sleep disruption is also believed to increase the risk of developing Alzheimer’s disease (AD), and likely plays a role in the progression of AD pathology and cognitive decline (Wang and Holtzman, 2019). Therefore, sleep is an important therapeutic target through all stages of life. Multiple recent studies using transcriptomics, proteomics, phosphoproteomics, or electron microscopy show that synapses of the mouse forebrain undergo widespread and profound remodeling during sleep, or in response to sleep deprivation (Bruning et al., 2019; de Vivo et al., 2017; Diering et al., 2017; Noya et al., 2019; Wang et al., 2018), strongly supporting synapses as a major locus for sleep’s restorative effects (Diekelmann and Born, 2010; Tononi and Cirelli, 2014). Sleep and synapses are highly dynamic during development. In animals and humans, sleep amount peaks at birth and declines sharply through childhood and adolescence. Sleep quality is also dynamic: Rapid-eye-movement (REM) sleep dominates early life while non-REM (NREM), also called slow-wave sleep (SWS), becomes more prominent during adolescence (Frank, 2015; Kurth et al., 2010; Nelson et al., 2013; Rensing et al., 2018; Wintler et al., 2020). Early postnatal life is dominated by synaptogenesis. During the remainder of childhood and adolescence, synapses are selectively pruned as neural circuits are refined (Penzes et al., 2011; Volk et al., 2015), and synapse density stabilizes during the transition to adulthood. During the period of synaptic pruning, elimination of synapses occurs predominantly during sleep (Li et al., 2017; Maret et al., 2011), and the adolescent peak in SWS is caused by the maturation of synapses in the cortex (Frank, 2015; Tononi and Cirelli, 2014). Treatment of sleep disruption throughout the lifespan will require further molecular insights into the synaptic signaling process activated during sleep at each life stage.

The endocannabinoid (eCB) system forms a prominent signaling pathway that regulates synaptic transmission throughout much of the brain (Castillo et al., 2012) and is known to play a role in the regulation of sleep, cognition, and other behaviors (Busquets-Garcia et al., 2011; Kesner and Lovinger, 2020). eCBs are bioactive lipids synthesized in post-synaptic compartments that signal in a retrograde manner by acting on pre-synaptic CB1 receptors to regulate neurotransmitter release (Di Marzo, 2018). The two most well-studied endocannabinoids, and known endogenous agonists for pre-synaptic CB1 receptors are 2-arachidonyl glycerol (2-AG) and anandamide (also called arachidonoyl ethanolamide, AEA) (Devane et al., 1992; Di Marzo, 2018; Mechoulam et al., 1995; Sugiura et al., 1995). Importantly, these two eCB metabolites have completely non-overlapping synthetic and degradative pathways, strongly suggesting that they are independently regulated and serve distinct functions. 2-AG is synthesized by the rate limiting enzyme diacylglycerol lipase alpha (DAGLα) and degraded by monoacylglycerol lipase (MAGL). AEA is part of a larger family of N-acylethanolamides (NAEs), that are synthesized by N-acyl phosphatidylethanolamine-specific phospholipase D, or through multiple secondary pathways. NAEs are degraded by a single lipase, fatty acid amide hydrolase (FAAH) (Di Marzo, 2018). The pharmaceutical industry and basic research have devoted considerable effort to the development of drugs that can act as selective agonists or antagonists for CB1 receptors as well as selectively upregulating 2-AG or NAEs through inhibition of MAGL or FAAH respectively (Ahn et al., 2009; Di Marzo, 2018; Niphakis et al., 2013).

NREM sleep involves highly coordinated neuronal activity throughout much of the brain, particularly the cortex. Cortical slow waves (0.5-4Hz) form the most prominent features in electroencephalogram (EEG) recordings during NREM sleep. eCB signaling has been shown to support the cortical up-state, a prominent mode of microcircuit activity within the cortex that underlies the generation of cortical slow waves (Pava et al., 2014; Steriade et al., 1993). Multiple studies have used pharmacology or genetics to target the eCB system in rodents and have shown that enhancement of eCB signaling promotes NREM stability, whereas inhibition of eCB signaling causes NREM fragmentation (Bogathy et al., 2019; Huitron-Resendiz et al., 2004; Kesner and Lovinger, 2020; Pava et al., 2016; Santucci et al., 1996). Thus, eCB signaling is well positioned to support the highly coordinated neuronal activity that likely promotes the restorative actions of NREM sleep. However, it is not known if eCB signaling promotes sleep in the context of development in both sexes.

In the current study, we combine quantitative mass spectrometry and behavioral pharmacology in mice to examine the role of eCB signaling in promoting sleep during development and in adulthood of both sexes. Our data show that the regulation of synaptic eCBs undergo a clear maturation where AEA and related NAEs are down-regulated during sleep deprivation in juveniles, transitioning to regulation of AEA and related metabolites by circadian mechanisms in adults. We confirm previous findings that enhancement of eCB signaling promotes sleep stability and suppression of eCB signaling drives sleep fragmentation in adult males (Bogathy et al., 2019; Pava et al., 2016), and we extend these findings to include developmental ages in both sexes. eCB signaling is important to sustain sleep in both males and females. However, we find that developing and adult males are far more sensitive to the sleep-promoting effects of eCB boosting drugs than females. Thus our findings further support a role for the eCB system in promoting sleep and uncover differences in this system during development vs. adulthood and between sexes.

## Methods

### Mice

All experiments were performed using C57Bl/6 mice purchased from Jackson Labs or Charles River, or bred in house. Unless otherwise specified, experiments were performed using juvenile (postnatal day P21-35), adolescent (P42-P56), or adult (>P90) mice. Additional experiments were performed using 8 month old females. All animal procedures were approved by the Institutional Animal Care and Use Committee of the University of North Carolina at Chapel Hill and were performed in accordance with the guidelines of the U.S. National Institutes of Health.

### Drugs and treatments

PF3845, JZL195, MJN110, were purchased from Cayman Chemicals. Drugs were dissolved in DMSO and then prepared into vehicle solution (5%DMSO, 5% Koliphor, 90% of a 1%NaCl solution) at the doses indicated. For metabolomics experiments, mice were treated with vehicle or the drugs indicated by intraperitoneal injection (IP) at Zeitgeber time 0 (ZT0) and then sacrificed 4 hours later at ZT4, followed by isolation of the forebrain and synaptosome preparation. For sleep behavior experiments, mice were injected immediately after the onset of the light phase (ZT0) or immediately preceding the onset of the dark phase (ZT12). Each mouse received an injection of vehicle or drug followed by a second injection 72 hours later of vehicle or drug with a cross over design. Sleep behavior was examined for 24 hours following each injection. The effects of the drug were determined by comparing 24hours following drug injection to the 24 hours following vehicle injection for each mouse.

### Synaptosome preparation

Male and female juvenile mice postnatal day 21 (P21), or young adults P56, were sacrificed during the sleep phase (ZT4), wake phase (ZT16), or following 4hours of total sleep deprivation (SD4) from ZT0-ZT4. Sleep deprivation was achieved using gentle handling, a low stress method that involves tapping on the mouse cage and disturbing the bedding material (Suzuki et al., 2013). Mouse forebrains, consisting of whole cortex and hippocampus, were dissected in ice-cold phosphate buffered saline, and then frozen on dry ice, and kept at -80°C until further processing. Frozen mouse forebrains were homogenized using 12 strokes from a glass homogenizer in ice-cold homogenization solution (320mM sucrose, 10mM HEPES pH 7.4, 1mM EDTA, 5mM Na pyrophosphate, 1mM Na_3_VO_4_, 200nM okadaic acid, 50nM JZL195, protease inhibitor cocktail (Roche)). Brain homogenate was then centrifuged at 1000xg for 10min at 4oC to obtain the P1 (nuclear) and S1 (post-nuclear) fractions. The S1 fraction was then layered on top of a discontinuous sucrose density gradient (0.8M, 1.0M or 1.2M sucrose in 10mM HEPES pH 7.4, 1mM EDTA, 5mM Na pyrophosphate, 1mM Na_3_VO_4_, 200nM okadaic acid, 50nM JZL195, protease inhibitor cocktail (Roche)) and then subjected to ultra-centrifugation at 82,500xg for 2hr at 4oC. Material accumulated at the interface of 1.0M and 1.2M sucrose (synaptosomes) was collected. Synaptosomes were diluted using 10mM HEPES pH7.4 (containing protease, phosphatase, and lipase inhibitors) to restore the sucrose concentration back to 320mM. The diluted synaptosomes were then pelleted by centrifugation at 100,000xg for 30min at 4oC. The synaptosome pellet was then resuspended in homogenization buffer (containing inhibitors). The protein concentration was determined using Bradford assay and material was frozen at -80oC prior to targeted mass spectrometry. Note the addition of dual MAGL/FAAH lipase inhibitor JZL195 50nM was added to all steps of sample preparation to maintain the integrity of eCB metabolites during sample preparation.

### Targeted mass spectrometry for endocannabinoid quantification

Frozen forebrain synaptosome fractions were prepared for endocannabinoid analysis as follows. Briefly, synaptosomes in homogenization buffer were removed from -80oC freezer and thawed on ice. The samples were then immediately placed into a microcentrifuge at 14,000RPM for 10 minutes at 4oC. The supernatant was removed and then 440μl of methanol, 50μl of internal standard containing 200ng/ml each of arachidonyl ethanolamide-d4 and Oleoyl ethanolamide-d4 and 2000ng/ml of 2-arachidonlyl glycerol-d5, and 10μl of 5mg/ml BHT in ethanol was added. The synaptosome pellet was resuspended and then vortexed for 5 seconds. The sample was then centrifuged at 14,000RPM for 10 minutes at 4oC. The supernatant was removed and then placed into a capped autosampler vial for analysis. LC/MS/MS analysis of endocannabinoids was performed as previously described (Gouveia-Figueira and Nording, 2015), with some modifications. Briefly, mass spectrometric analysis was performed on an Agilent 6490 triple quadrupole mass spectrometer in positive ionization mode. Calibration standards were analyzed over a range of concentrations from 0.2–40 pg on column for all of the ethanolamides. The following lipids were quantified: 2-arachidonyl glycerol (2-AG), arachidonoyl ethanolamide (AEA), docosahexaenoyl ethanolamide (DHEa), docosatetraenoyl ethanolamide (DEA), linoleoyl ethanolamide (LEA), oleoyl ethanolamide (OEA), palmitoleoyl ethanolamide (POEA), palmitoyl ethanolamide (PEA), stearoyl ethanolamide (SEA). Quantitation of endocannabinoids was performed using Agilent Masshunter Quantitative Analysis software. All results were normalized to protein concentration.

### Sleep phenotyping and behavior analysis

Mice were moved to our wake/sleep behavior satellite facility on a 12 hours:12 hours light:dark cycle (lights on 7am to 7pm). C57/BL-6 mice were individually housed in 15.5 cm^2^ cages with bedding, food and water. Mice were given two full dark cycles to acclimate to the recording cages before the beginning of data collection. No other animals were housed in the room during these experiments. Sleep and wake behavior were recorded using the PiezoSleep 2.0 monitoring system (Signal Solutions), which measures animal movement and breathing using piezoelectric sensors. 72 hours of uninterrupted data was collected for baseline behavior. Mice were recorded continuously before and after injections. Mechanical signals from the piezoelectric sensors are translated into electrical signals. Frequencies of mechanical activity is analyzed in sliding 2-second intervals centered at approximately 3 Hz, the typical breath rate for mice, to determine consistent respiration as sleep and inconsistent respiration as wake. Thresholds for sleep and wake were automatically calculated for each animal using statistical software (SleepStats, Signal Solutions). This method has been previously validated using simultaneous EEG recordings (Mang et al., 2014). Sleep amount and bout lengths were calculated for 24hrs following each injection, data separated into 12hr periods of light and dark.

### Statistical analysis

Microsoft Excel or Prism 8.0 (GraphPad Software Inc.) was used to analyze data from sleep behavior experiments outlined above. Paired two-tailed student’s t-test analyses were used to compare 24hrs sleep behavior, separated into 12hr light and dark periods, between vehicle injections and drug injections in the same animals. Comparisons of eCB metabolites between wake, sleep, and sleep deprivation conditions, or comparing vehicle treated groups to drug treatment was conducted using unpaired two-tail student’s t-test with Bonferroni correction for multiple comparisons. All data is presented as mean ± SEM, *p<0.05, **p<0.01, ***p<0.001.

## Results

### Sleep and circadian regulation of synaptic endocannabinoids

Daily fluctuations in endocannabinoids (eCBs) have been reported in blood and cerebral spinal fluid, from animal models and humans (Hillard, 2018; Murillo-Rodriguez et al., 2006; Valenti et al., 2004). However, it is not clear how these daily rhythms reflect the activity of eCBs at synapses, where eCBs are generated in post-synaptic neurons in response to neuronal activity (Castillo et al., 2012). Moreover, it is not clear how daily regulation of eCBs may differ during development compared to adulthood. Therefore, we developed a targeted mass spectrometry (MS) assay to quantify 2-AG, and NAEs, including AEA, from mouse forebrain synaptosome fractions. In preliminary assessment of this assay we quantified eCBs from whole forebrain homogenate, the S2 fraction (cytosol and extracellular fluid), and synaptosomes. We found that proportionally greater levels of eCBs were detected in synaptosomes compared to whole forebrain homogenate, and eCBs were essentially absent from the soluble S2 fraction (not shown), suggesting that brain eCBs are primarily localized to synaptic fractions and are retained during sample preparation. All steps of synaptosome isolation included the dual MAGL/FAAH inhibitor JZL195 (Long et al., 2009) to preserve the integrity of eCBs during sample preparation.

We then went on to examine the sleep-dependent regulation of synaptic eCBs during development and in adults. Male and female juvenile (P21) or young adult (P56) mice were sacrificed 4 hours into the sleep or wake cycle, ZT4 or ZT16 respectively, or following 4 hours of total sleep deprivation (SD4) using gentle handling starting from light onset (ZT0-ZT4) (Fig. 1A). Forebrains (hippocampus and cortex) were dissected followed by isolation of synaptosomes and targeted MS quantification. In juveniles we found that there was no difference in any of the eCBs compounds between wake and sleep, although AEA showed a trend to increase during sleep compared to wake. Compared to wake, SD4 caused a strong trend to decreased levels of AEA, and a significant decrease in every other species of NAE measured (Fig. 1B). These findings show that in juveniles, synaptic eCBs lack a daily rhythm but do show a clear response to sleep deprivation. In contrast, in young adults we found that AEA and OEA were significantly increased during sleep or SD4 (both collected at ZT4) compared to wake (Fig. 1C). All of the NAE species are FAAH substrates and showed their highest expression during sleep compared to wake or SD4. Consistent with this observation we recently showed using proteomics that synapse associated FAAH protein is decreased during sleep (Diering et al., 2017). 2-AG showed no changes between the three conditions at either age. Together these data show that as the mice mature towards adulthood, the eCB system transitions to regulation by the circadian rhythm, where synaptic AEA/OEA are increased during the sleep phase.

**Figure 1.**
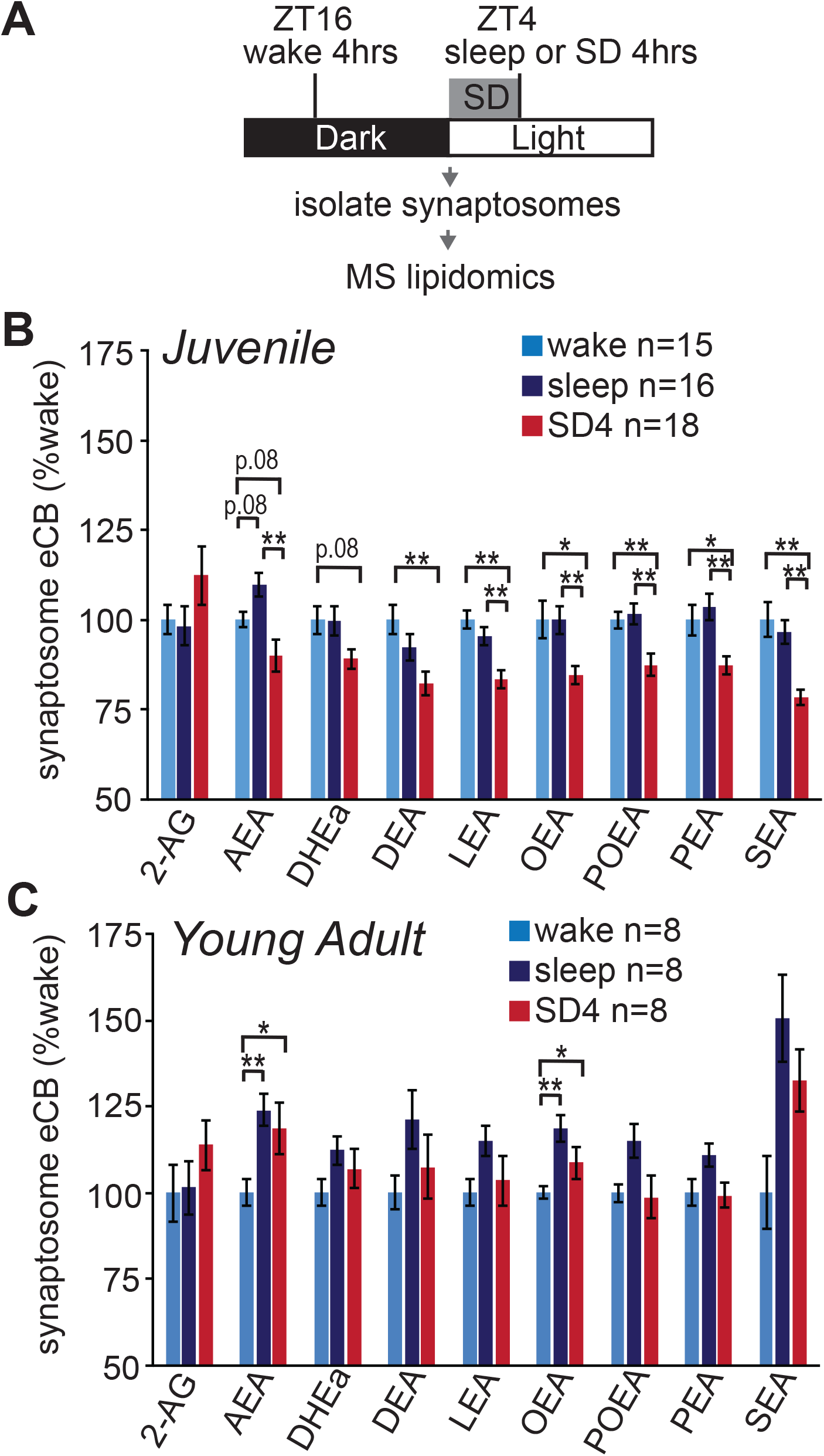
Regulation of synaptic eCBs during the sleep wake cycle in juvenile (P21) and young adult mice (P56). (**A**) Experimental design: Mice were maintained on 12:12hr light/dark schedule. Mice collected 4 hours into the wake period (ZT16) or sleep period (ZT4), or after 4 hours of gentle handling sleep deprivation (SD4) during the light period (ZT0-ZT4). The forebrain was dissected, the synaptosome fraction isolated, and quantified with mass spectrometry (MS). (**B**) Quantification of synaptic eCBs in juveniles. Data are normalized to the wake condition. n=15-18 per condition, includes males and females. (**C**) Quantification of synaptic eCBs in young adults. Data are normalized to the wake condition. n=8 per condition, includes males and females. Lipids quantified: 2-arachidonyl glycerol (2-AG), arachidonoyl ethanolamide (AEA), docosahexaenoyl ethanolamide (DHEa), docosatetraenoyl ethanolamide (DEA), linoleoyl ethanolamide (LEA), oleoyl ethanolamide (OEA), palmitoleoyl ethanolamide (POEA), palmitoyl ethanolamide (PEA), stearoyl ethanolamide (SEA). *P<0.05, **P<0.01 (unpaired two-tailed Student’s t-test with Bonferroni correction). Error bars indicate ± SEM. See also supplementary figure 1.

Note, results shown in Figure 1 were obtained from males and females in approximately equal numbers. We did not observe any significant differences or compelling trends in the levels of eCBs between the sexes at either age (not shown). Nonetheless, we doubled the number of samples for the juvenile age group in order to power an appropriate sex comparison. Highly similar results were obtained in juvenile males and females (supplementary Fig. 1). Thus we conclude that the levels of synaptic eCB metabolites and their regulation are not different between sexes.

Based on these findings, we hypothesized that pharmacological manipulation of eCBs using inhibitors of MAGL or FAAH would have differential sleep-promoting effects in juveniles vs. adults. Selective inhibitors MJN110 (5mg/kg) or PF3845 (10mg/kg) were used to block MAGL or FAAH respectively (Ahn et al., 2009; Niphakis et al., 2013). To confirm the action of these compounds, and to show that these treatments are able to manipulate synaptic eCBs, we treated adult mice at ZT0 by IP injection, followed by sacrifice at ZT4 and targeted quantification of eCBs in forebrain synaptosomes. As expected, compared to vehicle treated controls, FAAH inhibitor PF3845 caused no change in 2-AG and a 3-5 fold increase in NAE species (Fig. 2B). MAGL inhibitor MJN110 caused a 6-fold increase in 2-AG, but also an unexpected ∼30% decrease in AEA and the related NAEs (Fig. 2B). For comparison, treatment of dual MAGL/FAAH inhibitor JZL195 (10mg/kg) significantly increased the levels of 2-AG and all NAE species, consistent with a dual MAGL/FAAH inhibitor (Long et al., 2009). The decrease in NAE species following MJN110 treatment was not expected based on previous publications (Niphakis et al., 2013). An off-target effect of MJN110 acting on FAAH would be expected to result in increased NAEs, similarly to JZL195 (Fig. 2B), and so the observed decrease is likely to result from a secondary effect of elevated 2-AG. We went on to test the effects of MJN110 or PF3845 on sleep behavior, with the interpretation that MJN110 exerts effects through increased 2-AG and decreased NAEs, while PF3845 acts through an increase in NAEs with no change in 2-AG (Fig. 2B).

**Figure 2.**
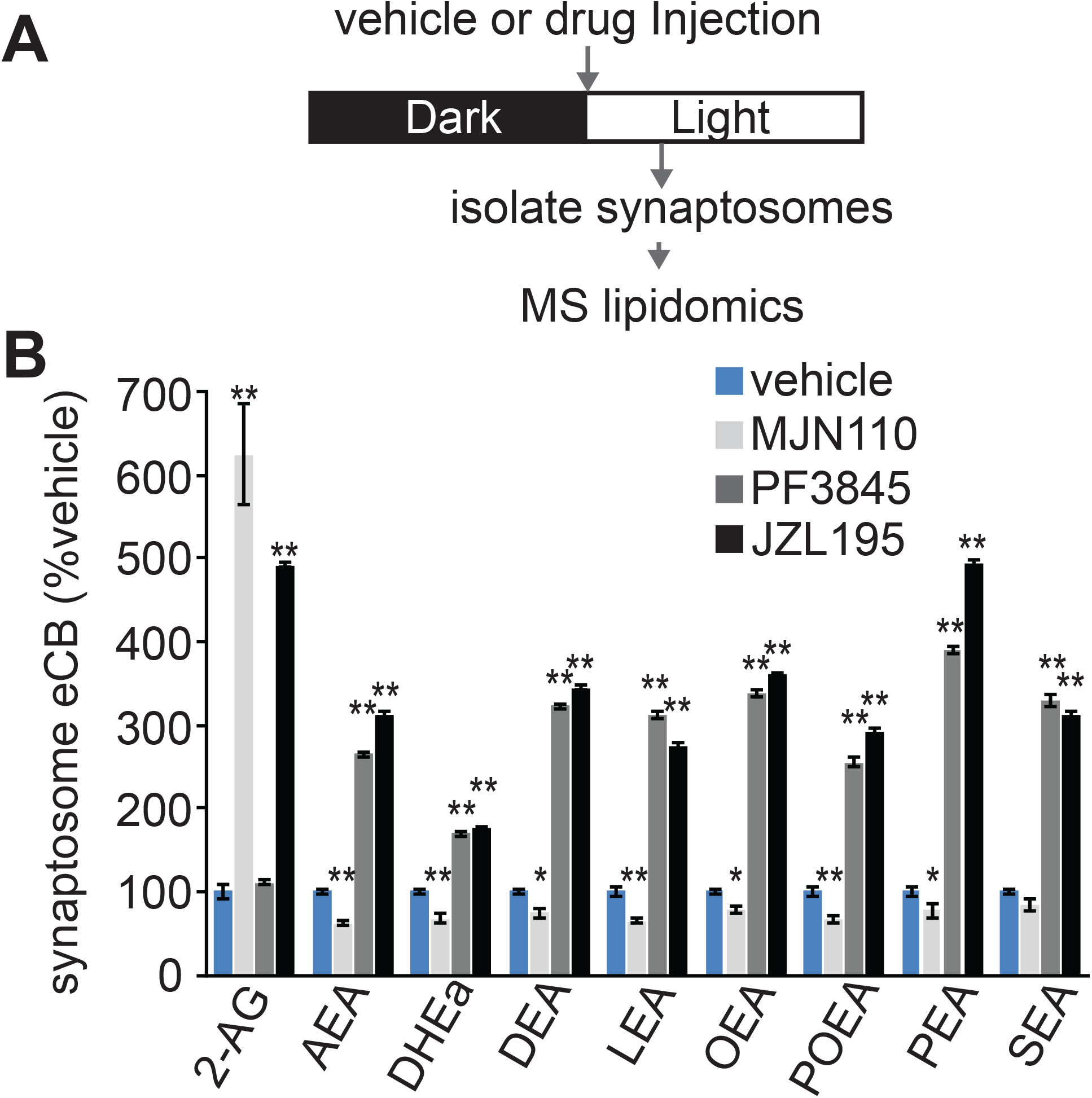
Pharmacological manipulations of synaptic eCB metabolites. (**A**) Experimental design: Mice were treated with vehicle or drugs with intraperitoneal injection at ZT0. Mice were sacrificed at ZT4, forebrain dissected, synaptosomes isolated and quantified with targeted MS analysis. (**B**) Quantification of synaptic eCBs in vehicle or drug treated adult mice. Data are normalized to the vehicle treatment condition. n=4-5 per condition. Treatment with MJN-110 (5mg/kg), PF3845 (10mg/kg), or JZL195 (5mg/kg). *P<0.05, **P<0.01 indicates significant difference between drug treatment and vehicle treatment control. Unpaired two-tailed Student’s t-test. Error bars indicate ± SEM.

### Sleep promoting effects of 2-AG or NAEs, during development and between sexes

We next went on to examine how pharmacological manipulations of eCB signaling affect sleep behavior in the context of development and between sexes. While we did not observe differences between the sexes in the levels of eCB metabolites, previous studies have shown sex-specific behavioral responses to pharmacological manipulation of the eCB system (Cooper and Craft, 2018). We measured sleep behavior in juvenile (P21), adolescent (P42), or adult (P100) male and female mice using a non-invasive piezo-electric home cage monitoring system, PiezoSleep, that uses highly sensitive piezoelectric polymers to measure mouse motion and breathing. This mechanical signal is then analyzed using custom software to score wake and sleep behavior. The PiezoSleep system has been previously validated using simultaneous EEG recordings (Mang et al., 2014). Note the ages indicated represent the age at the beginning of the experiment. Mice were given 2-3 days to acclimate to the recording cages prior to data collection. In the following experiments, male and female mice of indicated ages were treated by 2 consecutive IP injections of either vehicle or drug, separated by 3 days, using a random crossover design. Injections occurred either at ZT0 (lights on) or immediately prior to ZT12 (lights off) and sleep behavior was examined for 24hrs following injection (separated into 12hr blocks of light and dark). The effects of the drug were compared directly to the effects of vehicle control for each mouse.

Multiple previous studies have shown that increased eCB signaling by direct agonist treatment, or inhibition of MAGL or FAAH, promotes sleep in adult male mice or rats (Murillo-Rodriguez et al., 1998; Pava et al., 2016). Whether similar effects are also seen in developing mice or females has not been tested. Based on our quantification of synaptic eCBs in Fig. 1, our expectation was that increasing the levels of eCBs in males or females would have a more profound effect on sleep behavior in adults than in juveniles. We began by treating mice with MJN110 or PF3845 immediately prior to dark onset (ZT12), the beginning of the active phase, where sleep-promoting drugs are expected to show a more profound effect. In males, MJN110 treatment compared to vehicle control caused a significant increase in dark phase total sleep time and sleep bout length (Fig. 3A-C). Contrary to our expectations, qualitatively similar results were observed in male juveniles, adolescents, and adults. PF3845 treatment also significantly increased dark phase total sleep time and bout length in male juveniles, adolescents, and adults (Fig. 3D-F). The effects of either MJN110 or PF3845 were largely seen within the first 12hrs of treatment and had minimal effects on subsequent light phase sleep, with the exception of modest reduction in total sleep in some ages (Fig. 3B and E). In contrast, light phase injection of MJN110 or PF3845 in males had no significant effect on sleep amount or bout length at any age (supplementary Fig. 2). These findings show that in males, increased activity of 2-AG or NAEs in the dark phase can promote sleep and increase sleep bout length independent of developmental age. These data also confirm the sleep promoting effects of similar compounds tested in previous studies examining adult males (Pava et al., 2016).

**Figure 3.**
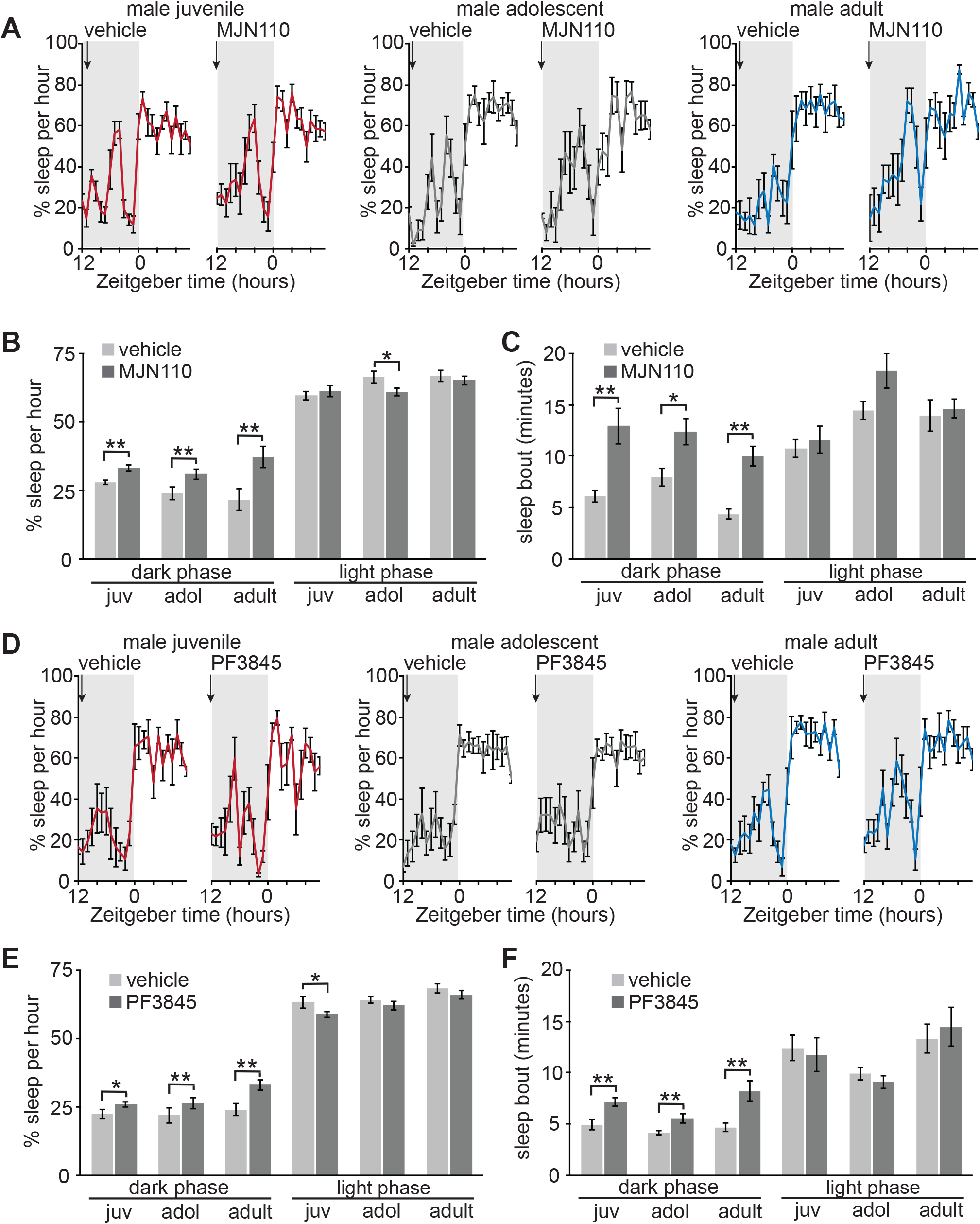
Increased 2-AG or AEA promotes dark phase sleep behavior in developing and adult males. (**A**) 24hr trace of average hourly sleep of male juveniles (juv, left), adolescents (adol, middle), and adults (right) treated with vehicle or MJN110 (5mg/kg) at the onset of the dark phase (ZT12), injection indicated by arrow. Grey bars in sleep traces indicate dark phase. (**B** and **C**) Quantification of average hourly sleep (**B**) and sleep bout length in minutes (**C**) for 24hrs following vehicle or MJN110 injection. Data separated into 12hrs of dark and light phases. N=12 juveniles, 8 adolescents, 8 adults. (**D**) 24hr trace of average hourly sleep of male juveniles (left), adolescents (middle) and adults (right) treated with PF3845 (10mg/kg) at the onset of the dark phase (ZT12), injection indicated by arrow. (**E** and **F**) Quantification of average hourly sleep (**E**) and sleep bout length in minutes (**F**) for 24hrs following vehicle or PF3845 injection. Data separated into 12hrs of dark and light phases. N=6 juveniles, 7 adolescents, 8 adults. *P<0.05 **P<0.001 (Paired two-tailed student’s t-test. Error bars indicate ± SEM. See also supplementary figure 2.

Identical dark-phase MJN110 or PF3845 treatments caused minimal effects in female mice, in striking contrast to males (Fig. 4). MJN110 treatment did not have any effect on dark phase total sleep in females at any age, but caused a significant increase in dark phase sleep bout length only in juvenile females. MJN110 also caused a modest decrease in subsequent light phase total sleep time in female adolescents (Fig. 4A-C). PF3845 showed no sleep promoting effects during the dark phase in females at any age and caused a decrease in subsequent light phase sleep bout length in female juveniles (Fig. 4D-F). Light phase injection of MJN110 caused a modest increase in light phase total sleep in female juveniles, but at the expense of reduced sleep bout length (supplementary Fig. 3). Light-phase injection of PF3845 in females had no measurable effect on sleep at any age (supplementary Fig. 3).

**Figure 4.**
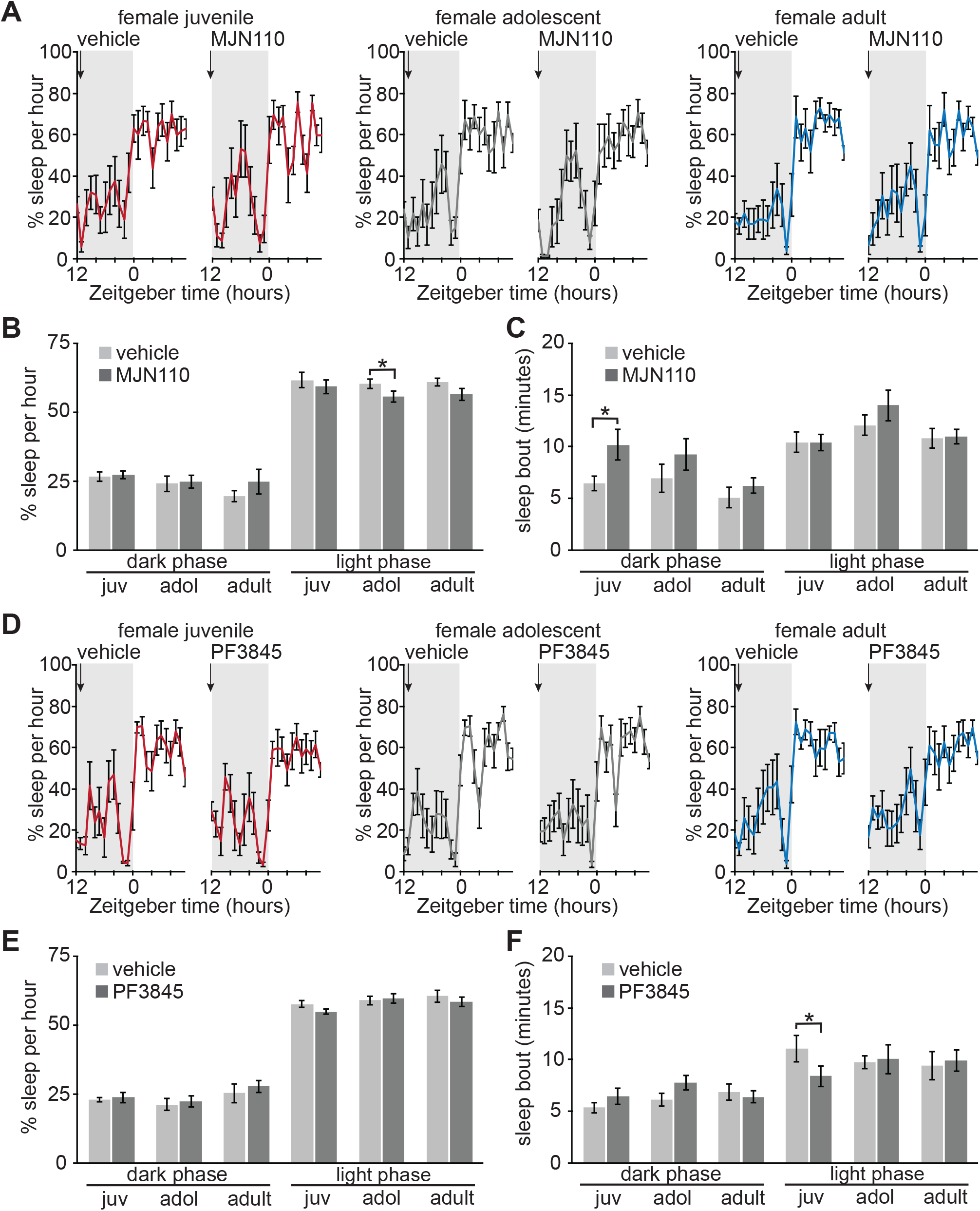
Increased 2-AG or AEA has minimal effects on dark phase sleep behavior in developing and adult females. (**A**) 24hr trace of average hourly sleep of female juveniles (left), adolescents (middle), and adults (right) treated with vehicle or MJN110 (5mg/kg) at the onset of the dark phase (ZT12), injection indicated by arrow. Grey bars in sleep traces indicate dark phase. (**B** and **C**) Quantification of average hourly sleep (**B**) and sleep bout length in minutes **(C)** for 24hrs following vehicle or MJN110 injection. Data separated into 12hrs of dark and light phases. N=8 juveniles, 8 adolescents, 8 adults. (**D**) 24hr trace of average hourly sleep of female juveniles (left), adolescents (middle) and adults (right) treated with PF3845 (10mg/kg) at the onset of the dark phase (ZT12), injection indicated by arrow. (**E** and **F**) Quantification of average hourly sleep (**E**) and sleep bout length in minutes (**F**) for 24hrs following vehicle or PF3845 injection. Data separated into 12hrs of dark and light phases. N=8 juveniles, 8 adolescents, 8 adults. *P<0.05 (Paired two-tailed Student’s t-test). Error bars indicate ± SEM. See also supplementary figure 3.

We had expected to see differential sleep promoting effects of MJN110 or PF3845 with respect to developmental age, but not sex. Instead we found that MJN110 or PF3845 treatments clearly promote sleep in males, independent of age, while females show no to minimal effects. Consistent with a previous report, the sleep-promoting effects of MAGL and FAAH inhibitor drugs in males are only observed in the dark phase (Pava et al., 2016).

### CB1 signaling sustains sleep in developing and adult males and females

Both males and females show an increase in AEA/OEA during the sleep phase as adults (Fig. 1), suggesting that eCBs do play a role in sleep in both sexes. However, pharmacological enhancement of 2-AG or NAEs, showed clear sleep promotion in males only. We hypothesize that an eCB tone supports sleep in both sexes, but this tone may be saturating in females, limiting the effects of MJN110 or PF3845, whereas in males the eCB tone may be unsaturated in the dark phase, allowing for the sleep promoting effects of MJN110 and PF3845. To test whether an eCB tone supports sleep in males and females, we examined the effects of treatment by a CB1 receptor inverse agonist, AM251 (10mg/kg) on sleep behavior. Previous studies (using males only) have shown mixed results of this, or similar treatments, in mice and rats. In some studies, CB1 inhibitors have no or modest effect (Calik and Carley, 2017; Goonawardena et al., 2011; Goonawardena et al., 2015), while in others these treatments suppress sleep behavior and cause sleep fragmentation (Bogathy et al., 2019; Pava et al., 2016). We suspected that these discrepancies may result from the different doses used. Therefore, we used AM251 at 10mg/kg, the higher end of the dose ranges reported in the literature. We first examined the effect of AM251 in the sleep (light) phase, when sleep disrupting compounds may be expected to show a more profound effect. Compared to vehicle control, light phase AM251 treatment caused a significant decrease in light phase total sleep time and sleep bout length, in juveniles and adults, both males and females (Fig. 5). The reduction in sleep bout length was particularly striking (Fig. 5C and F), suggesting that in the absence of CB1 signaling, sleep becomes highly fragmented, consistent with previous reports (Bogathy et al., 2019; Pava et al., 2016). In the accompanying sleep traces (Fig. 5A and D), the sleep suppressing effects of AM251 are particularly striking in the first hours of the light phase, probably reflecting the pharmacokinetics of this compound. The effects of AM251 were largely restricted to the first 12 hours (light phase), but sleep fragmentation was observed in the subsequent dark phase only in juvenile males. Adolescents were not tested in this experiment but our expectation based on these findings is that this age would respond similarly to juveniles and adults. Dark phase injection of AM251 in juvenile and adult males and females also caused a significant suppression of total sleep and sleep fragmentation within the first 12 hours (supplementary Fig. 4). Together these findings support our idea that an eCB tone supports sleep in both males and females, during development and adulthood, and that this eCB tone supports sleep in the light and dark phases.

**Figure 5.**
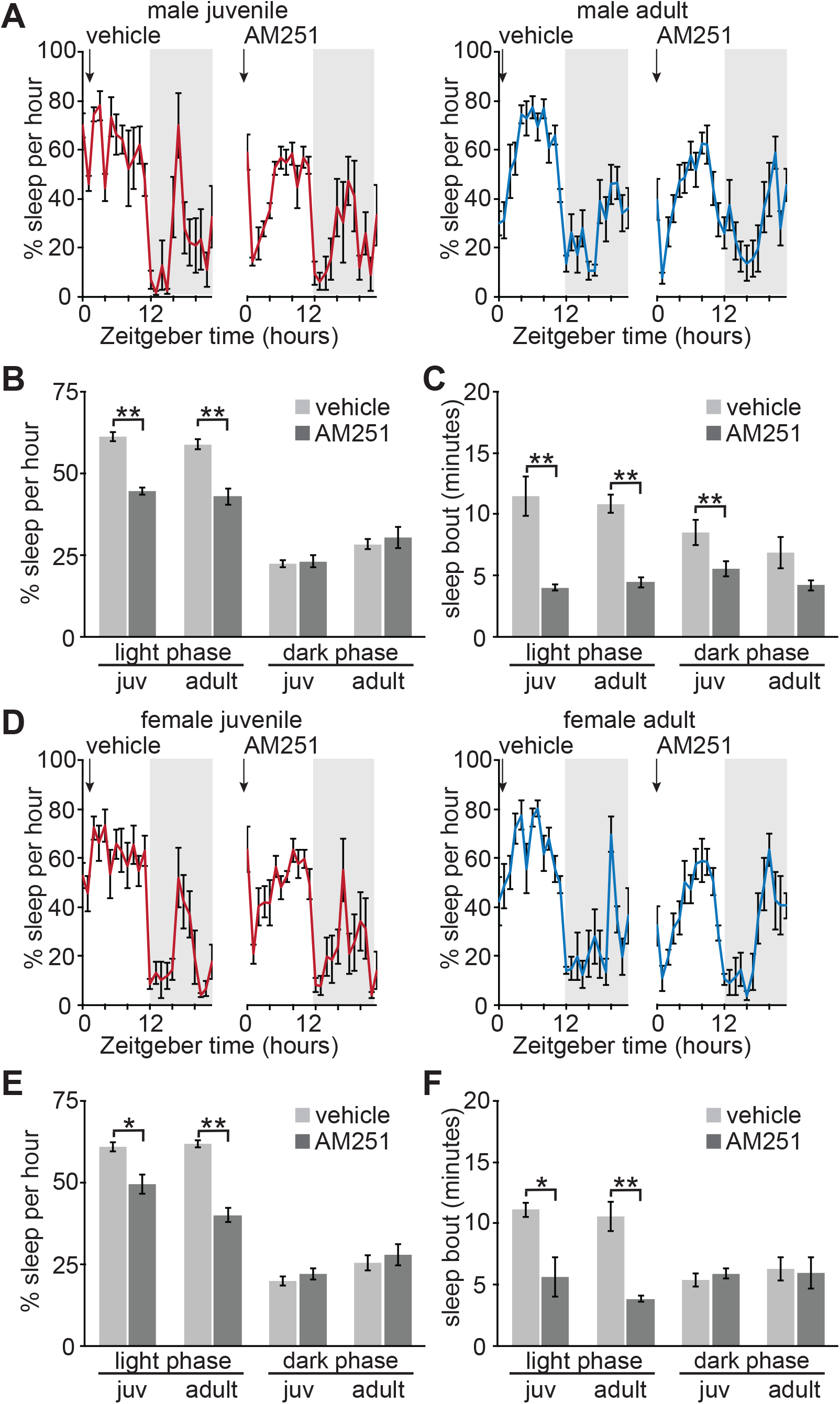
Tonic CB1 signaling sustains light phase sleep behavior in developing and adult males and females. (**A**) 24hr trace of average hourly sleep of male juveniles (left), and adults (right) treated with vehicle or AM251 (10mg/kg) at the onset of the light phase (ZT0), injection indicated by arrow. Grey bars in sleep traces indicate dark phase. (**B** and **C**) Quantification of average hourly sleep (**B**) and sleep bout length in minutes (**C**) for 24hrs following vehicle or AM251 injection. Data separated into 12hrs of light and dark phases. N=9 juveniles, 8 adults. (**D**) 24hr trace of average hourly sleep of female juveniles (left), and adults (right) treated with vehicle or AM251 (10mg/kg) at the onset of the light phase (ZT0). (**E** and **F**) Quantification of average hourly sleep (**E**) and sleep bout length in minutes (**F**) for 24hrs following vehicle or AM251 injection. Data separated into 12hrs of light and dark phases. N=8 juveniles, 7 adults. *P<0.05 **P<0.001 (Paired two-tailed Student’s t-test). Error bars indicate ± SEM. See also supplementary figure 4.

Returning to our idea that the eCB tone may be saturating in females, we tested the effects of MJN110 or AM251 in a group of older females, 8 months of age. Previous literature has suggested that eCB signaling may decrease with age (Bilkei-Gorzo, 2012; Di Marzo et al., 2015). We suspected that with further maturation, the eCB tone in females would decrease. Consistent with this idea, dark-phase injection of MJN110 caused a significant increase in total dark phase sleep and a trend of increased sleep bout length (supplementary Fig. 5A-C). This response is similar to what we observed in younger males, but not younger females. Light-phase injection of AM251 in older females caused a profound decrease in total sleep within the first 12hrs, and reduced sleep bout length that lasted 24hrs (supplementary Fig. 5D-F). Qualitatively, the sleep suppressing effects of AM251 appeared more severe in the 8month old females compared to 3months. Together these data suggest that an eCB tone supports sleep in females and that this tone declines with further maturation. Note that the 8 month old females tested here should not be considered representative of old age.

### Sleep-promotion by MJN110 or PF3845 require CB1

We wanted to confirm that increases in 2-AG or NAEs, via inhibition of MAGL or FAAH respectively, promote sleep by acting on CB1 receptors. This is particularly important for PF3845, because inhibition of FAAH increases synaptic levels all NAE metabolites (Fig. 2B) (Di Marzo, 2018), many of which are known to be bioactive. For example, FAAH substrates OEA and PEA are known to act as endogenous agonists for peroxisome proliferator-activated receptors (PPARs) nuclear transcription factors (Di Marzo, 2018). Adult male and female mice were injected prior to the dark phase with vehicle control or a cocktail containing AM251 (10mg/kg) together with either MJN110 (5mg/kg) or PF3845 (10mg/kg). Unlike MJN110 or PF3845 alone, which in males caused significant increases in total dark phase sleep and bout length, cocktail injections caused significant reductions, or a strong trend, in total sleep and bout length (Fig. 6A-H). Similar results were observed in males and females. In juvenile mice, treatment of AM251 with PF3845 also resulted in significantly reduced sleep time and bout length (supplementary Fig. 6). These results are highly similar to single treatment by AM251 and strongly suggest that the sleep promoting effects of MJN110 or PF3845 require CB1 as expected. The suppression of total sleep from cocktail injection were seen only in the first 12hrs (Fig. 6C and G), however, the reductions in sleep bout length were observed for 24hrs in both males and females (Fig. 6D and H).

**Figure 6.**
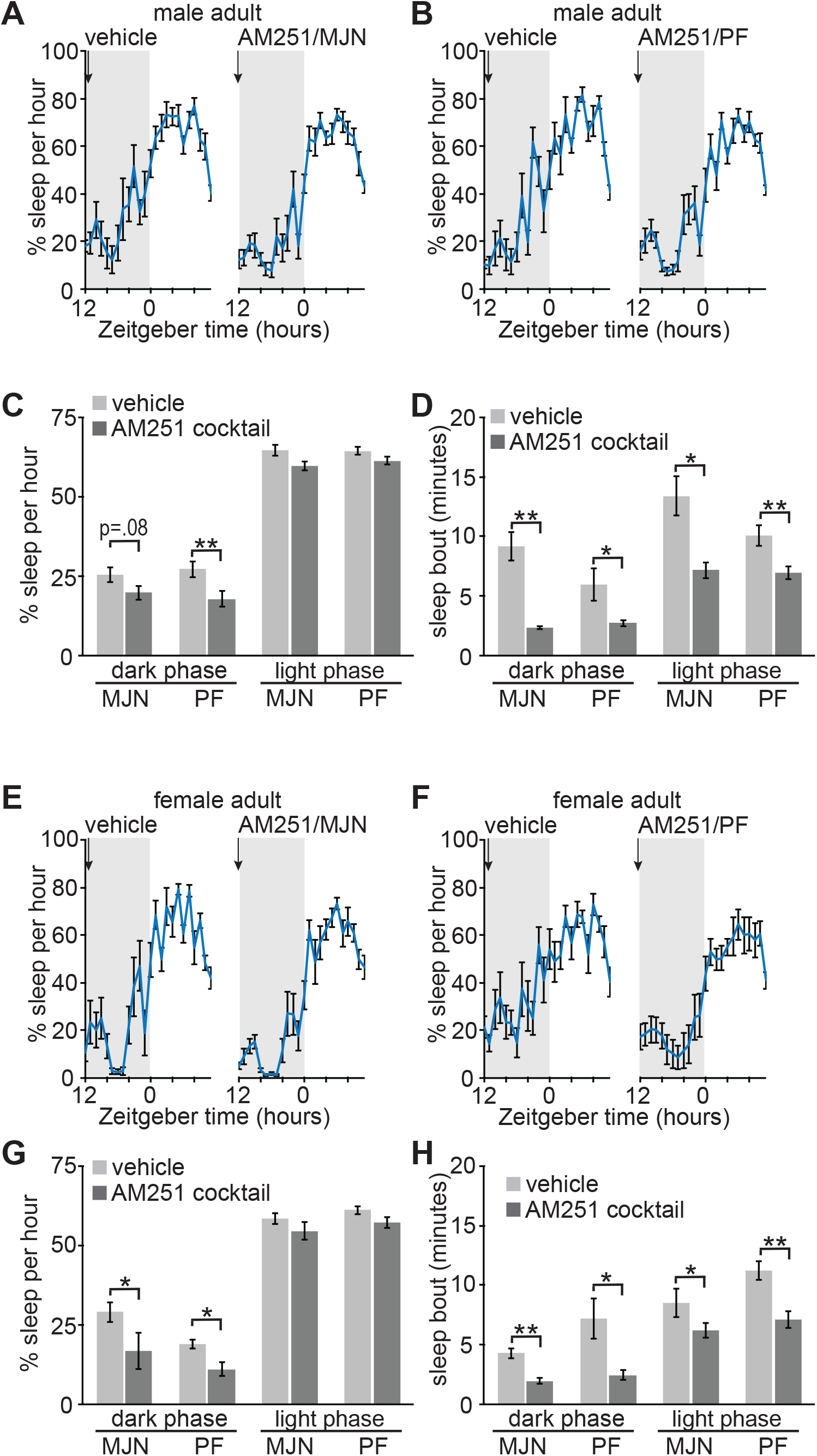
Sleep promoting effects of enhanced 2-AG or AEA in adults requires CB1. **(A** and **B**) 24hr trace of average hourly sleep of adult males treated with vehicle or combined AM251 (10mg/kg) / MJN110 (5mg/kg) (**A**), or AM251 (10mg/kg) / PF3845 (10mg/kg) (**B**). Injections, indicated by arrow, at the onset of the dark phase (ZT12). Grey bars in sleep traces indicate dark phase. (**C** and **D**) Quantification of average hourly sleep (**C**) and sleep bout length in minutes (**D**) for 24hrs following vehicle or combined drug injection. Data separated into 12hrs of dark and light phases. N=8 AM251/MJN110, 8 AM251/PF3845. (**E** and **F**) 24hr trace of average hourly sleep of adult females treated with vehicle or combined AM251 (10mg/kg) / MJN110 (5mg/kg) (**E**), or AM251 (10mg/kg) / PF3845 (10mg/kg) (**F**). (**G** and **H**) Quantification of average hourly sleep (**G**) and sleep bout length in minutes (**H**) for 24hrs following vehicle or combined drug injection. Data separated into 12hrs of dark and light phases. N=7 AM251/MJN110, 8 AM251/PF3845. *P<0.05 **P<0.001 (Paired two-tailed Student’s t-test). Error bars indicate ± SEM. See also supplementary figure 6.

## Discussion

In this study we used quantitative mass spectrometry and pharmacological manipulations targeting the eCB system to examine the role of eCB signaling in sleep behavior in the context of development. We systematically compared results between developing and adult mice and between males and females. We confirmed previous reports that enhancing the levels of 2-AG or AEA by inhibition of MAGL or FAAH respectively, enhanced sleep amount and bout length in adult males (Kesner and Lovinger, 2020; Pava et al., 2016), and we extend these findings by showing similar results in juvenile and adolescent males. In contrast, inhibition of MAGL or FAAH in developing and early adult females had minimal effects. Inhibition of CB1 with a direct antagonist, AM251, caused a clear suppression of sleep amount and sleep fragmentation in males and females, at all ages tested. Together, our findings suggest that an eCB tone is important to promote and sustain sleep in males and females, throughout development and into adulthood.

### Sex based differences in the eCB system

Suppression of CB1 signaling has a clear effect in both sexes, while the response to eCB boosting drugs show a clear male bias in our experiments. Quantification of synaptic eCBs by mass spectrometry did not indicate any sex differences in the levels of these metabolites in juveniles or young adults, suggesting that differences in response to MAGL/FAAH inhibition may result from differences in the expression levels or distribution of CB1 or downstream signaling components. A large body of literature shows clear sex differences in behavioral responses to cannabinoids or eCB targeted pharmacology (Cooper and Craft, 2018; Craft et al., 2013). Sex differences in the expression of CB1 mRNA or protein, or in the density of CB1 receptor binding using radio-labeled ligands, have been described in mice and rats but are highly region selective (Liu et al., 2020; Reich et al., 2009; Riebe et al., 2010). It is also plausible that sex-differences may exist in the signaling machinery downstream of CB1 activation. Expression of eCB system components is highly sensitive to pre-natal or early life drug exposures and stress in a brain region and sex-specific manner (Bara et al., 2018; Reich et al., 2009), further complicating our understanding of the sex-specific biology of this system. Sex-specific behavioral responses to eCB/CB1 targeted pharmacology is likely due to sexual dimorphism in specific cell populations and neural circuits. The neural circuits that underlie the sleep promoting effects of eCBs, particularly in males are not known, although the eCB system is most prominently expressed in the mouse forebrain, and CB1 mRNA is expressed at higher density in many regions of the male cortex (Liu et al., 2020). We hypothesize that CB1 expression may be limiting in females in the brain regions that underlie the sleep promoting effects of MJN110 or PF3845 treatment, while in males, higher CB1 expression permits this behavioral response. Thus, while the sexually divergent response to MJN110 and PF3845 we report here is in general agreement with the greater literature (Cooper and Craft, 2018; Craft et al., 2013), the underlying molecular, cellular, or circuit basis for this difference between sexes is not clear and will require further research.

### 2-AG vs. AEA

Increased signaling of CB1 by either 2-AG or AEA promoted sleep in males. Does it matter that there are two distinct eCB classes when either one can activate CB1 and promote sleep? We propose that CB1 signaling is required to promote sleep, but that the two major agonists are differentially regulated to serve distinct physiological functions. Our targeted quantification shows that synaptic eCBs of the NAE family undergo a clear maturation in their regulation. AEA and related NAEs are suppressed by sleep deprivation in juveniles, transitioning to circadian regulation of AEA/OEA in young adults, while 2-AG appears to be regulated independently of the sleep-wake cycle at both ages. 2-AG has been characterized as a “full-agonist” for CB1 receptors, but sustained increases in 2-AG through high doses of MAGL inhibitor drives the desensitization of CB1, leading to drug tolerance and secondary effects consistent with suppression of CB1 signaling (Busquets-Garcia et al., 2011; Di Marzo, 2018; Kinsey et al., 2013; Pava et al., 2016; Schlosburg et al., 2010; Sugiura et al., 1996). AEA is a “partial agonist” but does not drive desensitization, allowing for sustained CB1 signaling (Burkey et al., 1997; Schlosburg et al., 2010). Unlike other neurotransmitters that are stored in vesicles for regulated release, eCBs are synthesized from post-synaptic membranes, commonly described as synthesized “on demand” in response to neuronal activity. Prominent forms of eCB-dependent short-term plasticity including depolarization induced suppression of excitation or inhibition (DSE and DSI) fit this description. DSE and DSI have been shown to result from rapid and transient synthesis of 2-AG, but not AEA (Tanimura et al., 2010; Yoshino et al., 2011). A recent pre-print (https://doi.org/10.1101/2020.10.08.329169) describes the use of a novel eCB-biosensor called GRAB, and shows that phasic electrical stimulation of neurons results in rapid 2-AG synthesis, but not AEA. In other settings, eCB signaling is better described as being “tonic”, where an eCB tone likely due to AEA and not 2-AG, is constitutively acting on CB1 receptors over longer time scales (Gonzalez-Islas et al., 2012; Kim and Alger, 2010; Pava et al., 2014). We speculate that during the maturation of the cortex, a tonic mode of AEA signaling/metabolism regulated by the circadian rhythm emerges to promote light phase sleep activity in mice. Consistent with this, we previously used quantitative proteomics to examine how the synapse proteome is remodeled during the sleep/wake cycle in young adult mice and found that FAAH is downregulated at synapses during the sleep phase (Diering et al., 2017), matching our current dataset that FAAH substrates AEA and OEA are upregulated at synapses during the sleep phase (Fig. 1). We suggest that tonic AEA signaling, that does not drive desensitization of CB1, is better suited than 2-AG to support sustained signaling over the many hours of the sleep phase. In support of this, mice with deletion of FAAH have been shown to have sustained elevations in AEA and other NAEs, and to have increased total amount and bout lengths of NREM sleep, and higher levels of NREM delta power compared to wild type mice (Huitron-Resendiz et al., 2004).

### CB1 signaling, NREM sleep and development

Using our piezoelectric home cage recording apparatus, we are able to make accurate measurements of total sleep and bout length (Mang et al., 2014), but we are not able to make measurements of the proportions of sleep made up from REM and NREM sleep. Nonetheless, we believe the results of our experiments are most consistent with eCB-CB1 signaling playing a role in stabilizing NREM sleep. Treatment of male mice with MJN110 or PF3845 increased total dark phase sleep amount, but arguably had a greater effect on sleep bout length, a measurement of sleep stability. Similarly, inhibition of CB1 with AM251 in either sex reduced total sleep time, but additionally caused a very strong reduction in sleep bout length. Pava et al., indeed showed using EEG recording, that direct CB1 agonist treatment, or increased eCB levels, promoted NREM sleep and increased NREM sleep bout length, at the expense of wakefulness and REM sleep (Pava et al., 2016). As REM sleep can only be engaged from NREM, stabilization of NREM would be expected to reduce REM sleep through reduced NREM-REM transitions. Moreover, in an earlier publication, Pava et al., showed that tonic CB1 signaling is important to sustain the cortical up-state, a synchronized depolarization of cortical neurons that is a dominant mode of micro-circuit activity in the cortex during NREM sleep (Pava et al., 2014; Steriade et al., 1993). Synchronized Up-state-like activity highly reminiscent of NREM sleep is also observed in dissociated cortical cultures in vitro (Hinard et al., 2012; Kaufman et al., 2014; Saberi-Moghadam et al., 2018). In recent unpublished work we have observed that up-state like activity in primary cortical cultures also requires CB1 signaling, and that this tonic CB1 signaling is modulated by FAAH and AEA. The molecular mechanisms by which CB1 supports up-state activity and stabilizes NREM sleep is not known. One candidate downstream of CB1, is the activation of G-protein coupled inwardly rectifying potassium channels (GIRKs). GIRKs drive hyperpolarization potentially aiding in the synchronization of cortical neurons that underlies slow-wave activity seen in NREM. Indeed, a recent study showed that pharmacological activation of GIRK channels in mice promotes NREM sleep (Zou et al., 2019).

Sleep behavior is known to undergo an important maturation during the ages considered in this study. From weening at P21 to adulthood mice undergo a decrease in the levels of REM sleep transitioning to more consolidated episodes of NREM sleep (Nelson et al., 2013; Rensing et al., 2018). Matching this time course of maturation, we observe the emergence of a circadian regulation of AEA and related OEA. We speculate that the emergence of a circadian increase in the AEA tone during the sleep phase, stabilizes NREM episodes, thereby supporting the maturation of sleep behavior and the transition from REM dominant sleep to NREM. Sleep disruption, including frequent awakenings, is a major problem for many children with autism spectrum disorder (Mazzone et al., 2018; Missig et al., 2019). Unfortunately, sleep-aid medications for use in children are extremely limited. ASD is also diagnosed with a well-known male bias of ∼4:1, although the biological basis of this sex difference is not known. Declines in sleep amount and quality are also an expected component of aging. Our data show that increased eCB levels could promote sleep behavior in developing and adult male mice, and older adult females, suggesting that eCB based sleep-aids could be an effective treatment for sleep disruption at multiple life stages. However, the effect of such treatments on developmental and cognitive process will require further research.

Considerable interest is growing around the potential of the eCB system as a therapeutic target in a number of conditions. Our data support a role for the eCB system in promoting sleep, and we further extend this work to examine developmental time points. Our results also strongly emphasize that sex as a biological variable will require considerable attention in the ongoing development of eCB and cannabinoid based medicines.

## Acknowledgements

This work was supported by a grant to GHD from the National Institutes of Health (RF1AG068063).

## Figure legends

**Supplementary Figure 1.**
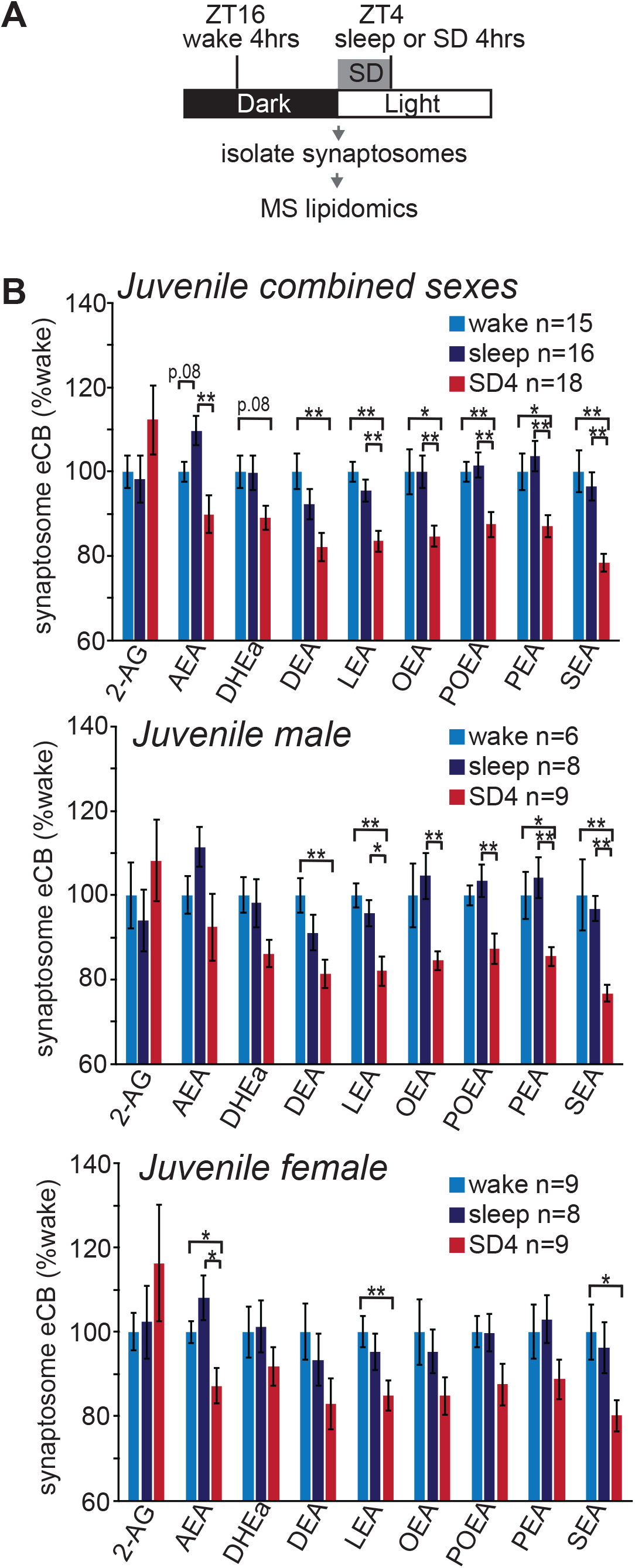
Similar regulation of eCBs during the sleep wake cycle between juvenile (P21) males and females. (**A**) Experimental design: Mice were maintained on 12:12hr light/dark schedule. Mice collected 4 hours into the wake period (ZT16) or sleep period (ZT4), or after 4 hours of gentle handling sleep deprivation (SD4) during the light period (ZT0-ZT4). The forebrain was dissected, the synaptosome fraction isolated, and quantified with mass spectrometry (MS). (**B**) Quantification of synaptic eCBs in juveniles: combined sexes; males only; females only. Data are normalized to the wake condition. N values per condition are indicated. Highly similar results are obtained in males or females. No significant differences were seen between males and females. *P<0.05, **P<0.01 (unpaired two-tailed Student’s t-test with Bonferroni correction). Error bars indicate ± SEM.

**Supplementary Figure 2.**
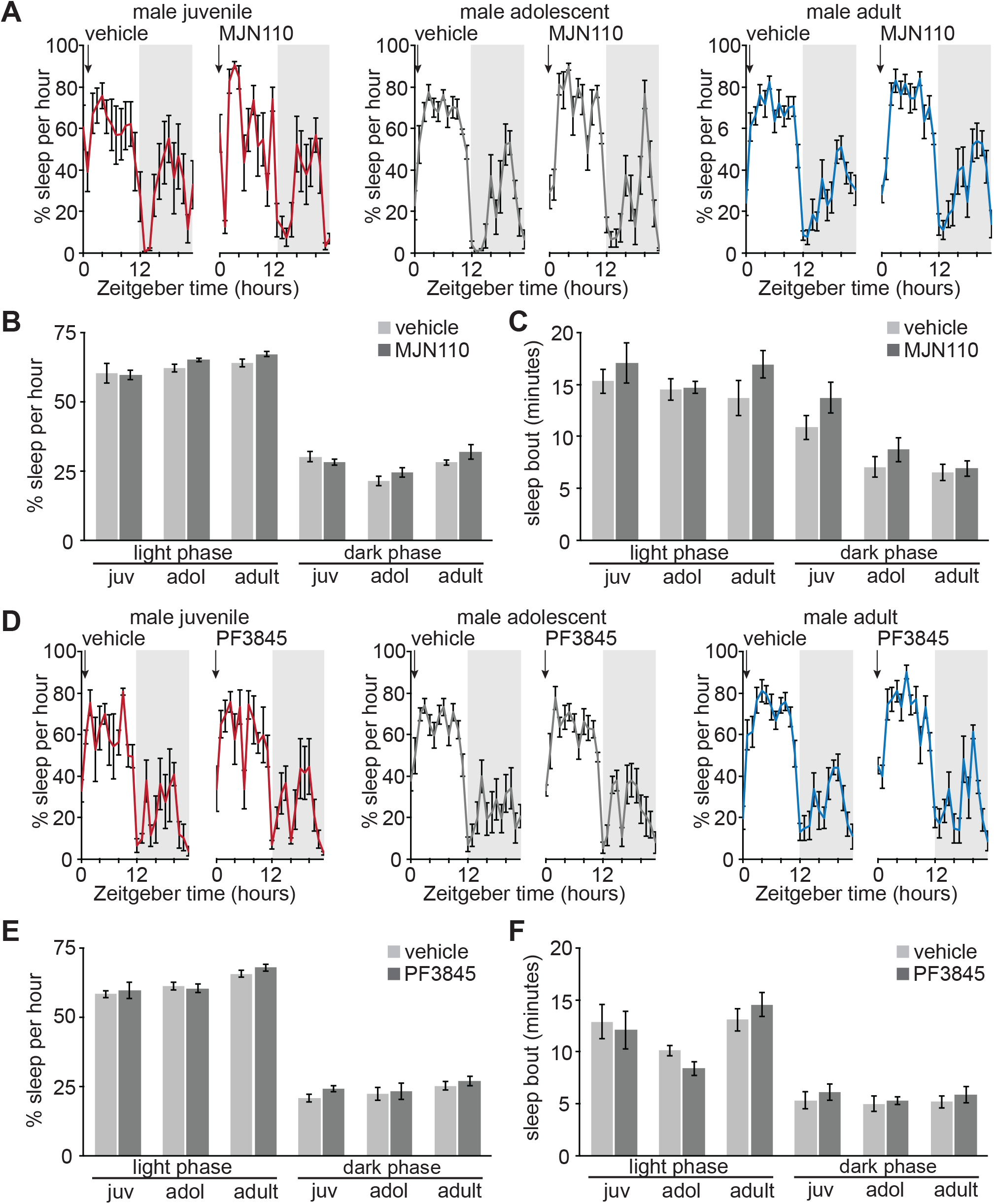
Increased 2-AG or AEA has no effect on light phase sleep behavior in developing and adult males. (**A**) 24hr trace of average hourly sleep of male juveniles (juv, left), adolescents (adol, middle), and adults (right) treated with vehicle or MJN110 (5mg/kg) at the onset of the light phase (ZT0), injection indicated by arrow. Grey bars in sleep traces indicate dark phase. (**B** and **C**) Quantification of average hourly sleep (**B**) and sleep bout length in minutes (**C**) for 24hrs following vehicle or MJN110 (MJN) injection. Data separated into 12hrs of light and dark phases. N=6 juveniles, 8 adolescents, 8 adults. (**D**) 24hr trace of average hourly sleep of male juveniles (left), adolescents (middle), and adults (right) treated with PF3845 (10mg/kg) at the onset of the light phase (ZT0), injection indicated by arrow. (**E** and **F**) Quantification of average hourly sleep (**E**) and sleep bout length in minutes (**F**) for 24hrs following vehicle or PF3845 injection. Data separated into 12hrs of light and dark phases. N=4 juveniles, 7 adolescents, 8 adults. *P<0.05 **P<0.001 (Paired two-tailed Student’s t-test). Error bars indicate ± SEM.

**Supplementary Figure 3.**
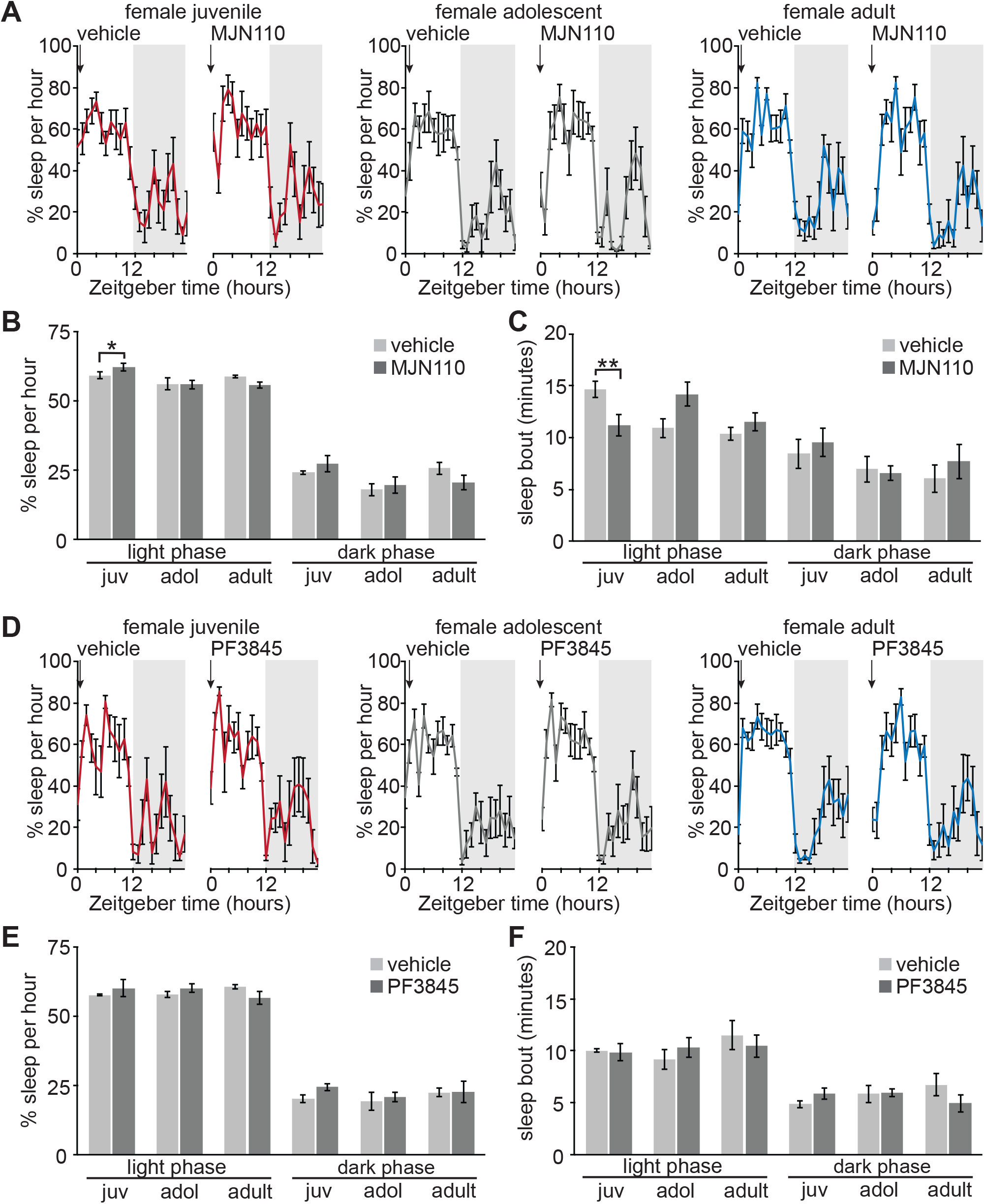
Increased 2-AG or AEA has modest effects on light phase sleep behavior in developing and adult females. (**A**) 24hr trace of average hourly sleep of female juveniles (left), adolescents (middle), and adults (right) treated with vehicle or MJN110 (5mg/kg) at the onset of the light phase (ZT0), injection indicated by arrow. Grey bars in sleep traces indicate dark phase. (**B** and **C**) Quantification of average hourly sleep (**B**) and sleep bout length in minutes (**C**) for 24hrs following vehicle or MJN110 injection. Data separated into 12hrs of light and dark phases. N=8 juveniles, 8 adolescents, 8 adults. (**D**) 24hr trace of average hourly sleep of female juveniles (left), adolescents (middle) and adults (right) treated with PF3845 (10mg/kg) at the onset of the light phase (ZT0), injection indicated by arrow. (**E** and **F**) Quantification of average hourly sleep (**E**) and sleep bout length in minutes (**F**) for 24hrs following vehicle or PF3845 injection. Data separated into 12hrs of light and dark phases. N=5 juveniles, 8 adolescents, 8 adults. *P<0.05, **P<0.01 (Paired two-tailed Student’s t-test). Error bars indicate ± SEM.

**Supplementary Figure 4.**
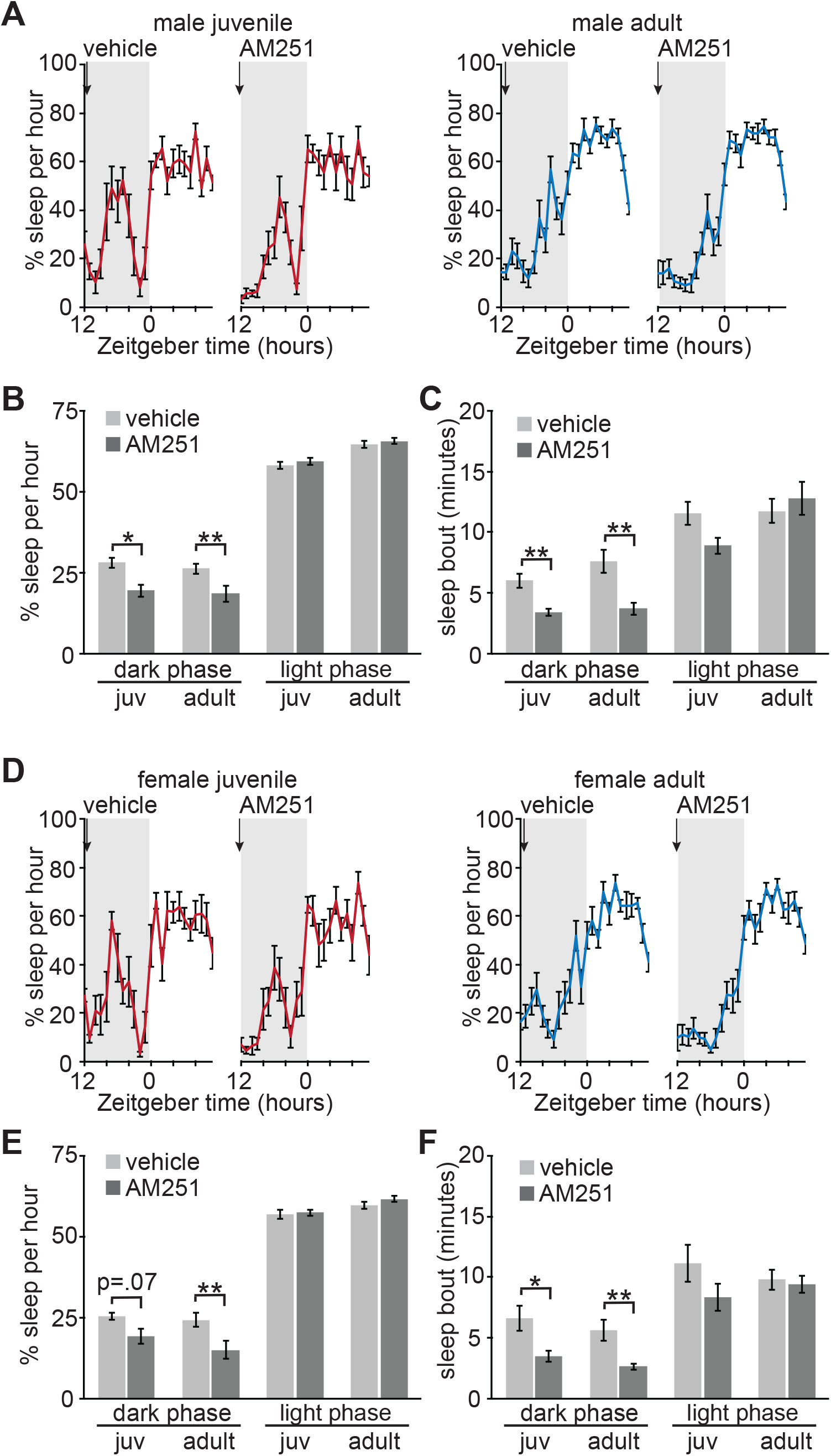
Tonic CB1 signaling sustains dark phase sleep behavior in developing and adult males and females. (**A**) 24hr trace of average hourly sleep of male juveniles (left), and adults (right) treated with vehicle or AM251 (10mg/kg) at the onset of the dark phase (ZT12), injection indicated by arrow. Grey bars in sleep traces indicate dark phase. (**B** and **C**) Quantification of average hourly sleep (**B**) and sleep bout length in minutes (**C**) for 24hrs following vehicle or AM251 injection. Data separated into 12hrs of dark and light phases. N=9 juveniles, 16 adults. (**D**) 24hr trace of average hourly sleep of female juveniles (left), and adults (right) treated with vehicle or AM251 (10mg/kg) at the onset of the dark phase (ZT12). (**E** and **F**) Quantification of average hourly sleep (**E**) and sleep bout length in minutes (**F**) for 24hrs following vehicle or AM251 injection. Data separated into 12hrs of dark and light phases. N=8 juveniles, 15 adults. *P<0.05 **P<0.001 (Paired two-tailed Student’s t-test). Error bars indicate ± SEM.

**Supplementary Figure 5.**
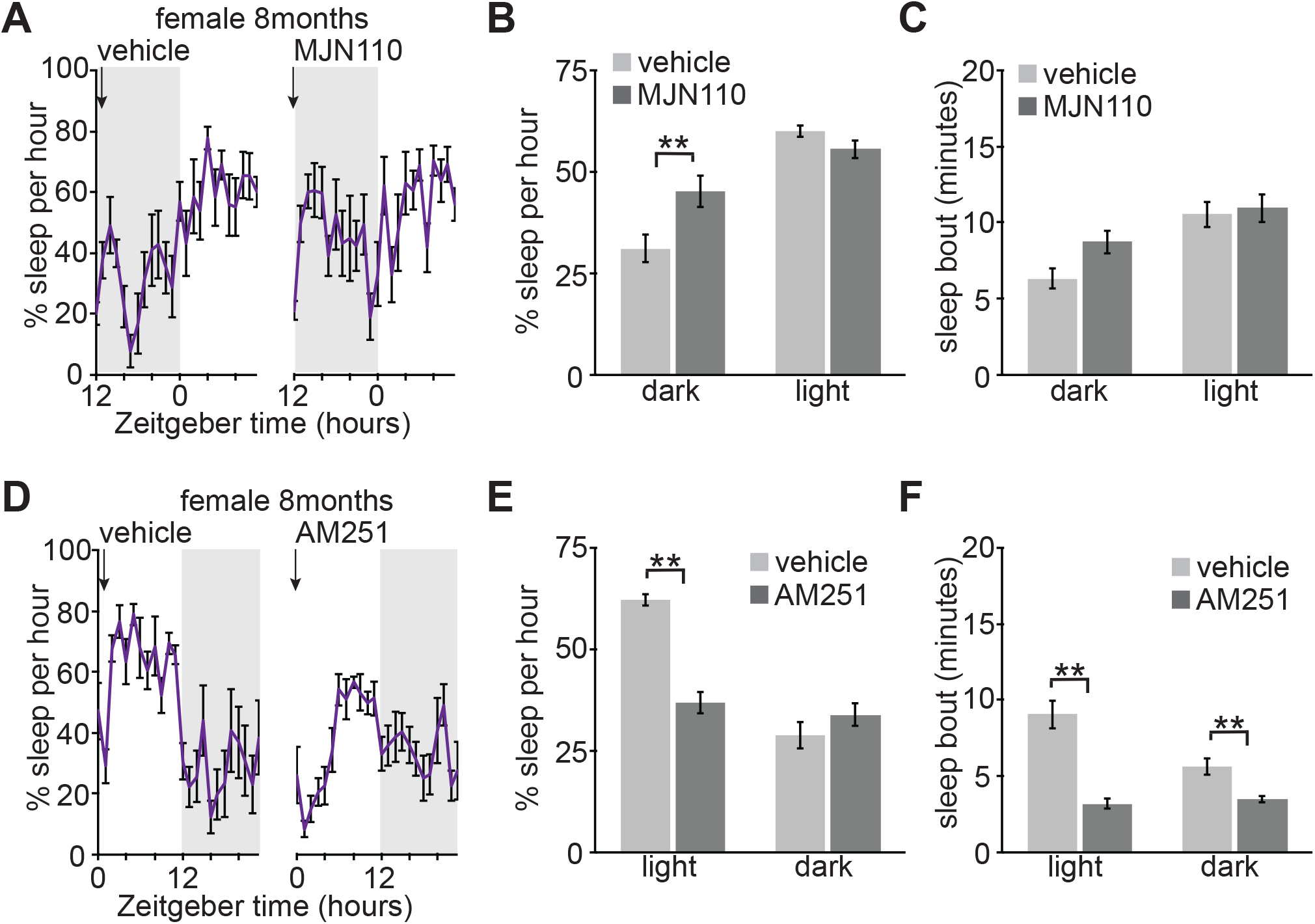
Mature 8 month old females respond to sleep promoting effects of increased 2-AG, and sleep suppressing effects of CB1 inhibition. (**A**) 24hr trace of average hourly sleep of 8month old females treated with vehicle or MJN110 (5mg/kg). Injections, indicated by arrow, at the onset of the dark phase (ZT12). Grey bars in sleep traces indicate dark phase. (**B** and **C**) Quantification of average hourly sleep (**B**) and sleep bout length in minutes (**C**) for 24hrs following vehicle or MJN110 injection. Data separated into 12hrs of dark and light phases. N=8. (**D**) 24hr trace of average hourly sleep of 8 month old females treated with vehicle or AM251 (10mg/kg). Injections, indicated by arrow, at the onset of the light phase (ZT0). (**E** and **F**) Quantification of average hourly sleep (**E**) and sleep bout length in minutes (**F**) for 24hrs following vehicle or AM251 injection. Data separated into 12hrs of light and dark phases. N=8. **P<0.001 (Paired two-tailed Student’s t-test). Error bars indicate ± SEM.

**Supplementary Figure 6.**
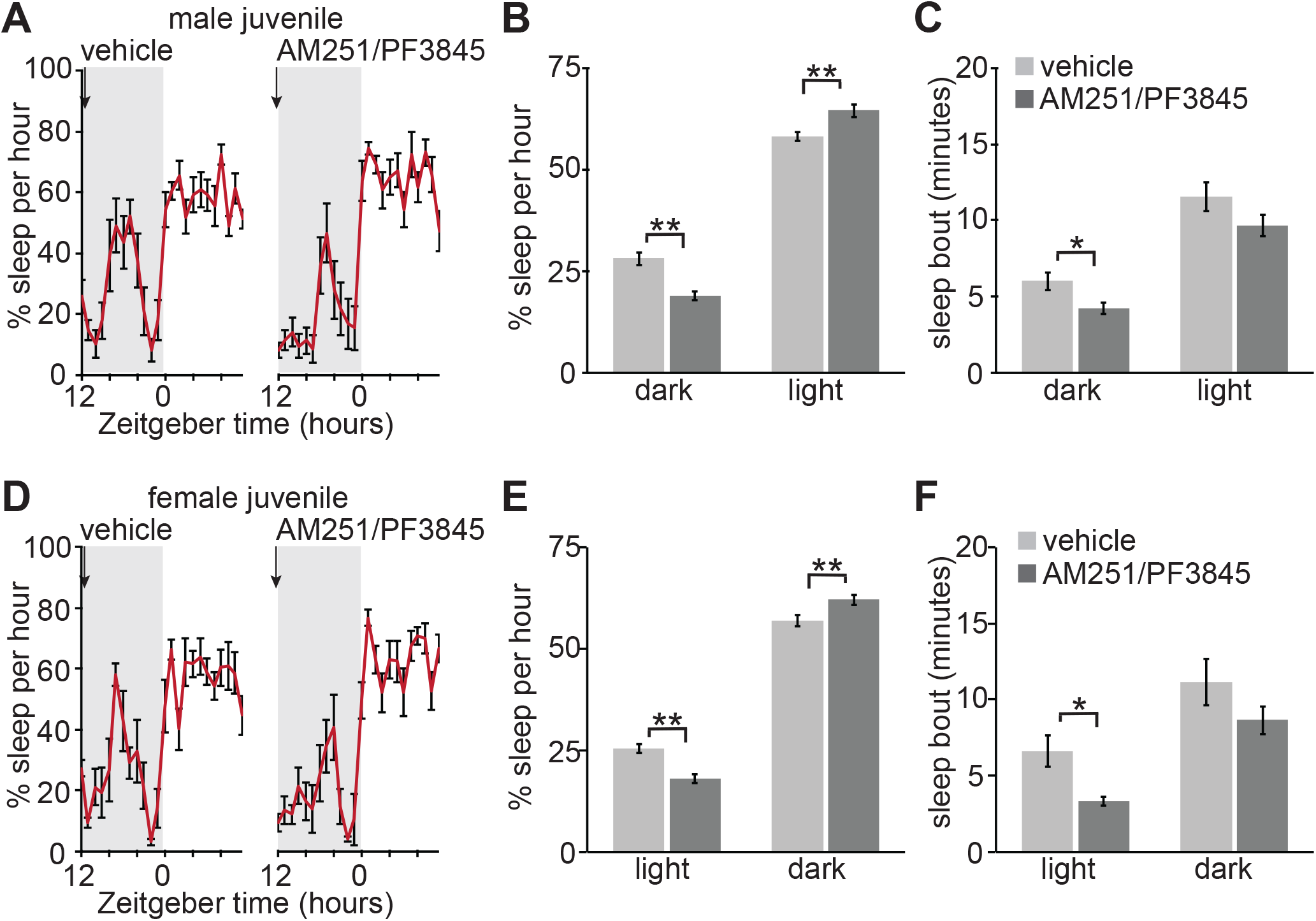
Sleep promoting effects of enhanced AEA in juveniles requires CB1. (**A**) 24hr trace of average hourly sleep of juvenile males treated with vehicle or combined AM251 (10mg/kg) / PF3845 (10mg/kg). Injections, indicated by arrow, at the onset of the dark phase (ZT12). Grey bars in sleep traces indicate dark phase. (**B** and **C**) Quantification of average hourly sleep (**B**) and sleep bout length in minutes (**C**) for 24hrs following vehicle or combined drug injection. Data separated into 12hrs of dark and light phases. N=9. (**D**) 24hr trace of average hourly sleep of juvenile females treated with vehicle or combined AM251 (10mg/kg) / PF3845 (10mg/kg). (**E** and **F**) Quantification of average hourly sleep (**E**) and sleep bout length in minutes (**F**) for 24hrs following vehicle or combined drug injection. Data separated into 12hrs of dark and light phases. N=8 *P<0.05 **P<0.001 (Paired two-tailed Student’s t-test). Error bars indicate ± SEM.

## References

Ahn, K., Johnson, D.S., Mileni, M., Beidler, D., Long, J.Z., McKinney, M.K., Weerapana, E., Sadagopan, N., Liimatta, M., Smith, S.E., et al. (2009). Discovery and characterization of a highly selective FAAH inhibitor that reduces inflammatory pain. Chem Biol 16, 411–420.

Bara, A., Manduca, A., Bernabeu, A., Borsoi, M., Serviado, M., Lassalle, O., Murphy, M., Wager-Miller, J., Mackie, K., Pelissier-Alicot, A.L., et al. (2018). Sex-dependent effects of in utero cannabinoid exposure on cortical function. Elife 7.

Bilkei-Gorzo, A. (2012). The endocannabinoid system in normal and pathological brain ageing. Philos Trans R Soc Lond B Biol Sci 367, 3326–3341.

Bogathy, E., Papp, N., Vas, S., Bagdy, G., and Tothfalusi, L. (2019). AM-251, A Cannabinoid Antagonist, Modifies the Dynamics of Sleep-Wake Cycles in Rats. Front Pharmacol 10, 831.

Bruning, F., Noya, S.B., Bange, T., Koutsouli, S., Rudolph, J.D., Tyagarajan, S.K., Cox, J., Mann, M., Brown, S.A., and Robles, M.S. (2019). Sleep-wake cycles drive daily dynamics of synaptic phosphorylation. Science 366.

Burkey, T.H., Quock, R.M., Consroe, P., Ehlert, F.J., Hosohata, Y., Roeske, W.R., and Yamamura, H.I. (1997). Relative efficacies of cannabinoid CB1 receptor agonists in the mouse brain. Eur J Pharmacol 336, 295–298.

Busquets-Garcia, A., Puighermanal, E., Pastor, A., de la Torre, R., Maldonado, R., and Ozaita, A. (2011). Differential role of anandamide and 2-arachidonoylglycerol in memory and anxiety-like responses. Biol Psychiatry 70, 479–486.

Calik, M.W., and Carley, D.W. (2017). Effects of Cannabinoid Agonists and Antagonists on Sleep and Breathing in Sprague-Dawley Rats. Sleep 40.

Castillo, P.E., Younts, T.J., Chavez, A.E., and Hashimotodani, Y. (2012). Endocannabinoid signaling and synaptic function. Neuron 76, 70–81.

Cooper, Z.D., and Craft, R.M. (2018). Sex-Dependent Effects of Cannabis and Cannabinoids: A Translational Perspective. Neuropsychopharmacology 43, 34–51.

Craft, R.M., Marusich, J.A., and Wiley, J.L. (2013). Sex differences in cannabinoid pharmacology: a reflection of differences in the endocannabinoid system? Life Sci 92, 476–481.

de Vivo, L., Bellesi, M., Marshall, W., Bushong, E.A., Ellisman, M.H., Tononi, G., and Cirelli, C. (2017). Ultrastructural evidence for synaptic scaling across the wake/sleep cycle. Science 355, 507–510.

Devane, W.A., Hanus, L., Breuer, A., Pertwee, R.G., Stevenson, L.A., Griffin, G., Gibson, D., Mandelbaum, A., Etinger, A., and Mechoulam, R. (1992). Isolation and structure of a brain constituent that binds to the cannabinoid receptor. Science 258, 1946–1949.

Di Marzo, V. (2018). New approaches and challenges to targeting the endocannabinoid system. Nat Rev Drug Discov 17, 623–639.

Di Marzo, V., Stella, N., and Zimmer, A. (2015). Endocannabinoid signalling and the deteriorating brain. Nat Rev Neurosci 16, 30–42.

Diekelmann, S., and Born, J. (2010). The memory function of sleep. Nat Rev Neurosci 11, 114–126.

Diering, G.H., Nirujogi, R.S., Roth, R.H., Worley, P.F., Pandey, A., and Huganir, R.L. (2017). Homer1a drives homeostatic scaling-down of excitatory synapses during sleep. Science 355, 511–515.

Frank, M.G. (2015). Sleep and synaptic plasticity in the developing and adult brain. Curr Top Behav Neurosci 25, 123–149.

Gonzalez-Islas, C., Garcia-Bereguiain, M.A., and Wenner, P. (2012). Tonic and transient endocannabinoid regulation of AMPAergic miniature postsynaptic currents and homeostatic plasticity in embryonic motor networks. J Neurosci 32, 13597–13607.

Goonawardena, A.V., Plano, A., Robinson, L., Platt, B., Hampson, R.E., and Riedel, G. (2011). A Pilot Study into the Effects of the CB1 Cannabinoid Receptor Agonist WIN55,212-2 or the Antagonist/Inverse Agonist AM251 on Sleep in Rats. Sleep Disord 2011, 178469.

Goonawardena, A.V., Plano, A., Robinson, L., Ross, R., Greig, I., Pertwee, R.G., Hampson, R.E., Platt, B., and Riedel, G. (2015). Modulation of food consumption and sleep-wake cycle in mice by the neutral CB1 antagonist ABD459. Behav Pharmacol 26, 289–303.

Gouveia-Figueira, S., and Nording, M.L. (2015). Validation of a tandem mass spectrometry method using combined extraction of 37 oxylipins and 14 endocannabinoid-related compounds including prostamides from biological matrices. Prostaglandins Other Lipid Mediat 121, 110–121.

Hillard, C.J. (2018). Circulating Endocannabinoids: From Whence Do They Come and Where are They Going? Neuropsychopharmacology 43, 155–172.

Hinard, V., Mikhail, C., Pradervand, S., Curie, T., Houtkooper, R.H., Auwerx, J., Franken, P., and Tafti, M. (2012). Key electrophysiological, molecular, and metabolic signatures of sleep and wakefulness revealed in primary cortical cultures. J Neurosci 32, 12506–12517.

Huitron-Resendiz, S., Sanchez-Alavez, M., Wills, D.N., Cravatt, B.F., and Henriksen, S.J. (2004). Characterization of the sleep-wake patterns in mice lacking fatty acid amide hydrolase. Sleep 27, 857–865.

Kaufman, M., Reinartz, S., and Ziv, N.E. (2014). Adaptation to prolonged neuromodulation in cortical cultures: an invariable return to network synchrony. BMC Biol 12, 83.

Kesner, A.J., and Lovinger, D.M. (2020). Cannabinoids, Endocannabinoids and Sleep. Front Mol Neurosci 13, 125.

Kim, J., and Alger, B.E. (2010). Reduction in endocannabinoid tone is a homeostatic mechanism for specific inhibitory synapses. Nat Neurosci 13, 592–600.

Kinsey, S.G., Wise, L.E., Ramesh, D., Abdullah, R., Selley, D.E., Cravatt, B.F., and Lichtman, A.H. (2013). Repeated low-dose administration of the monoacylglycerol lipase inhibitor JZL184 retains cannabinoid receptor type 1-mediated antinociceptive and gastroprotective effects. J Pharmacol Exp Ther 345, 492–501.

Kurth, S., Ringli, M., Geiger, A., LeBourgeois, M., Jenni, O.G., and Huber, R. (2010). Mapping of cortical activity in the first two decades of life: a high-density sleep electroencephalogram study. J Neurosci 30, 13211–13219.

Li, W., Ma, L., Yang, G., and Gan, W.B. (2017). REM sleep selectively prunes and maintains new synapses in development and learning. Nat Neurosci 20, 427–437.

Liu, X., Li, X., Zhao, G., Wang, F., and Wang, L. (2020). Sexual dimorphic distribution of cannabinoid 1 receptor mRNA in adult C57BL/6J mice. J Comp Neurol 528, 1986–1999.

Long, J.Z., Nomura, D.K., Vann, R.E., Walentiny, D.M., Booker, L., Jin, X., Burston, J.J., Sim-Selley, L.J., Lichtman, A.H., Wiley, J.L., et al. (2009). Dual blockade of FAAH and MAGL identifies behavioral processes regulated by endocannabinoid crosstalk in vivo. Proc Natl Acad Sci U S A 106, 20270–20275.

Mang, G.M., Nicod, J., Emmenegger, Y., Donohue, K.D., O’Hara, B.F., and Franken, P. (2014). Evaluation of a piezoelectric system as an alternative to electroencephalogram/ electromyogram recordings in mouse sleep studies. Sleep 37, 1383–1392.

Maret, S., Faraguna, U., Nelson, A.B., Cirelli, C., and Tononi, G. (2011). Sleep and waking modulate spine turnover in the adolescent mouse cortex. Nat Neurosci 14, 1418–1420.

Mazzone, L., Postorino, V., Siracusano, M., Riccioni, A., and Curatolo, P. (2018). The Relationship between Sleep Problems, Neurobiological Alterations, Core Symptoms of Autism Spectrum Disorder, and Psychiatric Comorbidities. J Clin Med 7.

Mechoulam, R., Ben-Shabat, S., Hanus, L., Ligumsky, M., Kaminski, N.E., Schatz, A.R., Gopher, A., Almog, S., Martin, B.R., Compton, D.R., et al. (1995). Identification of an endogenous 2-monoglyceride, present in canine gut, that binds to cannabinoid receptors. Biochem Pharmacol 50, 83–90.

Missig, G., McDougle, C.J., and Carlezon, W.A., Jr. (2019). Sleep as a translationally-relevant endpoint in studies of autism spectrum disorder (ASD). Neuropsychopharmacology.

Murillo-Rodriguez, E., Desarnaud, F., and Prospero-Garcia, O. (2006). Diurnal variation of arachidonoylethanolamine, palmitoylethanolamide and oleoylethanolamide in the brain of the rat. Life Sci 79, 30–37.

Murillo-Rodriguez, E., Sanchez-Alavez, M., Navarro, L., Martinez-Gonzalez, D., Drucker-Colin, R., and Prospero-Garcia, O. (1998). Anandamide modulates sleep and memory in rats. Brain Res 812, 270–274.

Nelson, A.B., Faraguna, U., Zoltan, J.T., Tononi, G., and Cirelli, C. (2013). Sleep patterns and homeostatic mechanisms in adolescent mice. Brain Sci 3, 318–343.

Niphakis, M.J., Cognetta, A.B., 3rd, Chang, J.W., Buczynski, M.W., Parsons, L.H., Byrne, F., Burston, J.J., Chapman, V., and Cravatt, B.F. (2013). Evaluation of NHS carbamates as a potent and selective class of endocannabinoid hydrolase inhibitors. ACS Chem Neurosci 4, 1322–1332.

Noya, S.B., Colameo, D., Bruning, F., Spinnler, A., Mircsof, D., Opitz, L., Mann, M., Tyagarajan, S.K., Robles, M.S., and Brown, S.A. (2019). The forebrain synaptic transcriptome is organized by clocks but its proteome is driven by sleep. Science 366.

Pava, M.J., den Hartog, C.R., Blanco-Centurion, C., Shiromani, P.J., and Woodward, J.J. (2014). Endocannabinoid modulation of cortical up-states and NREM sleep. PLoS One 9, e88672.

Pava, M.J., Makriyannis, A., and Lovinger, D.M. (2016). Endocannabinoid Signaling Regulates Sleep Stability. PLoS One 11, e0152473.

Penzes, P., Cahill, M.E., Jones, K.A., VanLeeuwen, J.E., and Woolfrey, K.M. (2011). Dendritic spine pathology in neuropsychiatric disorders. Nat Neurosci 14, 285–293.

Reich, C.G., Taylor, M.E., and McCarthy, M.M. (2009). Differential effects of chronic unpredictable stress on hippocampal CB1 receptors in male and female rats. Behav Brain Res 203, 264–269.

Rensing, N., Moy, B., Friedman, J.L., Galindo, R., and Wong, M. (2018). Longitudinal analysis of developmental changes in electroencephalography patterns and sleep-wake states of the neonatal mouse. PLoS One 13, e0207031.

Riebe, C.J., Hill, M.N., Lee, T.T., Hillard, C.J., and Gorzalka, B.B. (2010). Estrogenic regulation of limbic cannabinoid receptor binding. Psychoneuroendocrinology 35, 1265–1269.

Saberi-Moghadam, S., Simi, A., Setareh, H., Mikhail, C., and Tafti, M. (2018). In vitro Cortical Network Firing is Homeostatically Regulated: A Model for Sleep Regulation. Sci Rep 8, 6297.

Santucci, V., Storme, J.J., Soubrie, P., and Le Fur, G. (1996). Arousal-enhancing properties of the CB1 cannabinoid receptor antagonist SR 141716A in rats as assessed by electroencephalographic spectral and sleep-waking cycle analysis. Life Sci 58, PL103–110.

Schlosburg, J.E., Blankman, J.L., Long, J.Z., Nomura, D.K., Pan, B., Kinsey, S.G., Nguyen, P.T., Ramesh, D., Booker, L., Burston, J.J., et al. (2010). Chronic monoacylglycerol lipase blockade causes functional antagonism of the endocannabinoid system. Nat Neurosci 13, 1113–1119.

Steriade, M., Nunez, A., and Amzica, F. (1993). Intracellular analysis of relations between the slow (< 1 Hz) neocortical oscillation and other sleep rhythms of the electroencephalogram. J Neurosci 13, 3266–3283.

Sugiura, T., Kodaka, T., Kondo, S., Tonegawa, T., Nakane, S., Kishimoto, S., Yamashita, A., and Waku, K. (1996). 2-Arachidonoylglycerol, a putative endogenous cannabinoid receptor ligand, induces rapid, transient elevation of intracellular free Ca2+ in neuroblastoma x glioma hybrid NG108-15 cells. Biochem Biophys Res Commun 229, 58–64.

Sugiura, T., Kondo, S., Sukagawa, A., Nakane, S., Shinoda, A., Itoh, K., Yamashita, A., and Waku, K. (1995). 2-Arachidonoylglycerol: a possible endogenous cannabinoid receptor ligand in brain. Biochem Biophys Res Commun 215, 89–97.

Suzuki, A., Sinton, C.M., Greene, R.W., and Yanagisawa, M. (2013). Behavioral and biochemical dissociation of arousal and homeostatic sleep need influenced by prior wakeful experience in mice. Proc Natl Acad Sci U S A 110, 10288–10293.

Tanimura, A., Yamazaki, M., Hashimotodani, Y., Uchigashima, M., Kawata, S., Abe, M., Kita, Y., Hashimoto, K., Shimizu, T., Watanabe, M., et al. (2010). The endocannabinoid 2-arachidonoylglycerol produced by diacylglycerol lipase alpha mediates retrograde suppression of synaptic transmission. Neuron 65, 320–327.

Tononi, G., and Cirelli, C. (2014). Sleep and the price of plasticity: from synaptic and cellular homeostasis to memory consolidation and integration. Neuron 81, 12–34.

Valenti, M., Vigano, D., Casico, M.G., Rubino, T., Steardo, L., Parolaro, D., and Di Marzo, V. (2004). Differential diurnal variations of anandamide and 2-arachidonoyl-glycerol levels in rat brain. Cell Mol Life Sci 61, 945–950.

Volk, L., Chiu, S.L., Sharma, K., and Huganir, R.L. (2015). Glutamate synapses in human cognitive disorders. Annu Rev Neurosci 38, 127–149.

Wang, C., and Holtzman, D.M. (2019). Bidirectional relationship between sleep and Alzheimer’s disease: role of amyloid, tau, and other factors. Neuropsychopharmacology.

Wang, Z., Ma, J., Miyoshi, C., Li, Y., Sato, M., Ogawa, Y., Lou, T., Ma, C., Gao, X., Lee, C., et al. (2018). Quantitative phosphoproteomic analysis of the molecular substrates of sleep need. Nature 558, 435–439.

Wintler, T., Schoch, H., Frank, M.G., and Peixoto, L. (2020). Sleep, brain development, and autism spectrum disorders: Insights from animal models. J Neurosci Res 98, 1137–1149.

Yoshino, H., Miyamae, T., Hansen, G., Zambrowicz, B., Flynn, M., Pedicord, D., Blat, Y., Westphal, R.S., Zaczek, R., Lewis, D.A., et al. (2011). Postsynaptic diacylglycerol lipase mediates retrograde endocannabinoid suppression of inhibition in mouse prefrontal cortex. J Physiol 589, 4857–4884.

Zou, B., Cao, W.S., Guan, Z., Xiao, K., Pascual, C., Xie, J., Zhang, J., Xie, J., Kayser, F., Lindsley, C.W., et al. (2019). Direct activation of G-protein-gated inward rectifying K+ channels promotes nonrapid eye movement sleep. Sleep 42.

